# Distinct binding modes shape exploitable asymmetries between echinocandin and triterpenoid resistance landscapes in Fks1

**DOI:** 10.64898/2026.11.10.765593

**Authors:** Hans Carolus, Nagène Wang, Stef Jacobs, Alexandre K. Dubé, Dimitrios Sofras, Celia Lobo Romero, Romain Durand, Bram Spruijtenburg, Eelco F. J. Meijer, Patrick Van Dijck, Patrick Lagüe, Christian R. Landry

## Abstract

Echinocandins are the first-line treatment for invasive fungal infections and work by inhibiting the fungal β-1,3-glucan synthase Fks1. Recently, a first-in-class triterpenoid Fks1 inhibitor, ibrexafungerp, was approved for clinical use. The extent to which the difference between these drug classes alters the landscape of resistance mutations in Fks1 is unknown. We used deep mutational scanning (DMS) to quantify the effects of all single-amino-acid variants spanning 159 residues across the three canonical Fks1 resistance hotspots and flanking regions in *Saccharomyces cerevisiae*. Drug selections revealed widespread position-specific cross-resistance and differential resistance, including an ibrexafungerp-specific resistance cluster downstream of hotspot 2 that defines a novel triterpenoid-specific resistance hotspot. We defined this as hotspot 4 and redefined the boundaries of the remaining three canonical echinocandin hotspots, based on the mutational landscape of our DMS analyses. Susceptibility testing of individual *FKS1* variants of *S. cerevisiae* and *Candidozyma auris* strains confirmed conserved class-specific and cross-resistance patterns. Structure-guided ligand placement, molecular dynamics-based simulated annealing, and DMS-informed pose selection linked drug-specific phenotypes to distinct drug-target interaction networks, providing a molecular rationale for the observed differential resistance patterns. These data provide a framework for predicting antifungal resistance, informing treatment strategies and future drug design.

**One-Sentence Summary:** Fks1 resistance landscapes redefine hotspot boundaries and show both overlapping and distinctive resistance patterns between echinocandins and ibrexafungerp which are diagnostically and therapeutically important.

## Main Text

Invasive fungal infections are increasingly difficult to treat because of the limited antifungal arsenal and increasing incidence of antifungal drug resistance. The fungal cell wall offers a unique therapeutic target. Its major structural polymer, β-1,3-glucan, is synthesized by Fks1, the membrane-embedded catalytic subunit of the β-1,3-glucan synthase complex. High-resolution structures of Fks1 located the catalytic center at the membrane-cytoplasm interface and revealed a transmembrane channel for glucan extrusion, providing a framework for understanding enzyme function and drug–target interactions^1^.

Echinocandins (ECNs) are amphiphilic lipopeptides that noncompetitively inhibit Fks1. Currently, echinocandins are recommended as first-line therapy for invasive candidiasis, the predominant hospital-associated fungal infection and a major cause of morbidity and mortality^2^. They consist of caspofungin (CAS), micafungin (MCF), anidulafungin (AND), and the latest approved echinocandin, rezafungin (RZF)^3^. The main mechanism of resistance to echinocandins is target alteration, through substitutions concentrated in mutational hotspots: hotspot 1 (HS1) and hotspot 2 (HS2). A third, more elusive hotspot (HS3) was identified through site-directed mutagenesis in *Saccharomyces cerevisiae*^4^ and later shown to play a role in echinocandin resistance of the emergent pathogen *Candidozyma auris*^5,6^. *Recent cryo-EM work has clarified how CAS engages the Fks1–glucan complex, providing a structural basis for echinocandin inhibition*^7^. *Yet, how substitutions in the hotspot regions disrupt this interaction remains largely unresolved. Durand et al*. systematically mapped all amino-acid substitutions within the three hotspots and showed that these regions converge around a membrane-facing cavity containing both shared and compound-specific resistance determinants^8^. However, because this analysis was restricted to the historically defined hotspots, the full extent of the resistance landscape, the precise boundaries of each hotspot, and the residue-level mechanisms linking mutation to impaired drug engagement remain unclear.

The recent approval of the oral triterpenoid ibrexafungerp (IBX) brought a new antifungal drug class into clinical use for the first time in more than two decades^9^. Although triterpenoids and echinocandins both inhibit Fks1, their distinct chemical scaffolds suggest different drug-target interactions and resistance routes. Recent structures show that the triterpenoid enfumafungin binds at the extracellular membrane leaflet, engaging the surface formed by transmembrane helices 5 and 6 (TM5–TM6) through distributed hydrophobic interactions and remodeling of the surrounding lipid environment; these studies propose that triterpenoids inhibit Fks1 by stabilizing a nonproductive conformational state rather than by occupying the catalytic site^10,11^. This binding mode appears to differ from that of the echinocandin complex, which suggests that resistance to this new class could have distinct molecular bases. A small set of *FKS* mutations has been shown to reduce triterpenoid susceptibility *in vitro*^10-14^ but therapy-emergent triterpenoid resistance has not yet been reported in clinical isolates, likely reflecting its recent introduction and still-limited clinical exposure. It remains unclear where echinocandin and triterpenoid resistance landscapes overlap, where they diverge, and whether class-specific resistance can preserve therapeutic options, which could be guided by knowing the genotypes of the infecting strains.

Here, we used deep mutational scanning (DMS) to quantify the effects of all possible single-amino-acid substitutions across 159 residues (3,021 substitutions) spanning the canonical Fks1 hotspots and their flanking regions, covering the proposed binding sites of echinocandins and triterpenoids. We screened the resulting libraries against all licensed echinocandins and IBX, validated selected variants in *S. cerevisiae* and *C. auris*, and integrated the resulting mutational landscapes with DMS-informed structural modeling and simulated annealing refinement.

### Extension of echinocandin hotspot boundaries and identification of HS4

We targeted three 53-residue windows in Fks1 encompassing hotspot 1 (residues 604–656), hotspot 3 (residues 659–711), and hotspot 2 (residues 1328–1380) as shown in **Figure 1A**. Each target region was first replaced independently with the hygromycin (HYG) resistance cassette, generating three region-specific recipient strains. Using CRISPR-Cas9 technology, these strains were subsequently co-transformed with a pCas plasmid expressing a guide RNA targeting the HYG cassette and PCR-amplified synthetic oligonucleotide pools (oPools) containing the complete set of codon variants as donor DNA. Each HS library contained 6,784 variants – 53 positions × 32 NNK codons × four synonymous barcode markers – yielding 20,352 unique designed sequences across the three regions. The complete library was constructed twice independently, producing six pools of approximately 13,000 to 69,000 transformant colonies that were pooled per library and screened separately. Each pool was screened in a no-drug control and against CAS, MCF, AND, RZF, or IBX at concentrations producing approximately 50% growth-rate inhibition of the wild-type, as determined from dose–response measurements (**Supplementary Figure S1**). Approximately 1.5×10^6^ cells were inoculated in 10 mL culture volumes per condition and propagated for nine generations with serial dilution to the starting inoculum cell concentration every three generations. Allele frequencies were measured in the input populations before selection (t_0_) and after three passages (nine generations; t_3_). Genomic DNA was extracted from the t_0_ and t_3_ populations, and each mutagenized region was amplified by PCR. The resulting amplicons were indexed and subjected to deep sequencing. Demultiplexed reads were processed using the gyōza pipeline^15^ to assign reads to the expected codon–barcode sequences and quantify their frequencies in each population, as schematically shown in **Figure 1B**. Selection coefficients (*S*_v_) were calculated from changes in variant frequencies between t_0_ and t_3_, while retaining the four synonymous barcode estimates and the two independently constructed pool estimates for replication quality analyses (**Supplementary Figure S2**). Replication quality analyses confirmed that all synonymous barcode classes were retained after demultiplexing and that selection coefficients were reproducible across independently constructed libraries for most drug-selected conditions, particularly in HS1 and HS2. Lower correlations were mainly observed for no-drug controls and selected HS3 conditions, consistent with weaker directional selection and lower signal-to-noise, and are discussed in detail in **Supplementary Results, Annex 1**.

**Figure 1.**
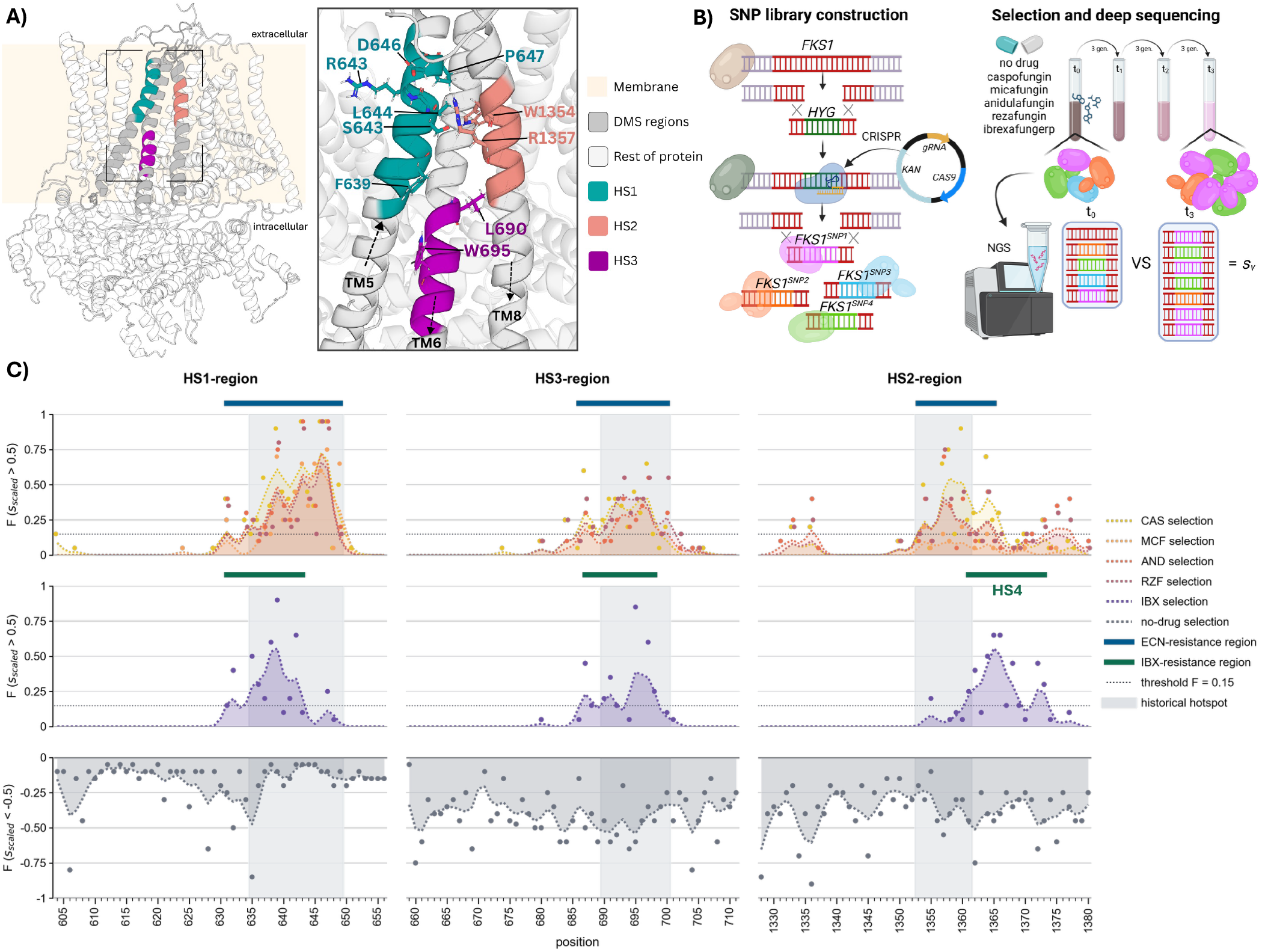
Deep mutational scanning of the Fks1 drug binding region and identification of resistance regions. **A)** Cryo-EM structure of *S. cerevisiae* Fks1 positioned within a modeled membrane, with the three DMS-targeted regions indicated on the protein structure. Historical echinocandin hotspot regions are highlighted within these targeted regions: HS1, E635-R649; HS3, L690-N700; and HS2, D1353-S1361. The zoomed structural view shows the directionality of the local transmembrane segments and the 10 most frequently reported echinocandin-resistance-associated residues according to FungAMR^16^, mapped to the corresponding *S. cerevisiae* Fks1 residues, summarized in **Supplementary Table S1**. Residues are colored according to their corresponding hotspot region. **B)** Schematic overview of the *FKS1* deep mutational scanning strategy. Variant libraries were introduced by CRISPR-Cas9-mediated targeting of the HYG marker that replaced the target region. Independently constructed libraries were selected for 9 generations in the absence of drug or in the presence of echinocandins or IBX. The input and selected populations were deep sequenced, and changes in variant frequencies between t_0_ and t_3_ were used to calculate generation-normalized selection coefficients (*S*_*v*_). **C)** Position-level resistance profiles across the three DMS-targeted *FKS1* regions. For each amino acid position and condition, nested mean selection coefficients were centered relative to the wild-type amino acid at the same position and scaled within each drug-region combination. Drug panels show the fraction of substitutions with strong positive scaled activity (*S*_scaled_ > 0.*5*). The no-drug panel shows the fraction of substitutions with strong negative scaled activity (*S*_scaled_ < −0.*5*). Points indicate unsmoothed position-level fractions and dotted lines with shaded areas show Gaussian-smoothed profiles. The dotted horizontal line indicates the resistance-region calling threshold of *F* = 0.15. Grey shaded areas indicate historical echinocandin hotspot intervals. Blue bars indicate DMS-defined ECN-resistance regions based on CAS, AND, and RZF; green bars indicate DMS-defined IBX-resistance regions. Resistance regions were defined as clusters of positions meeting the activity criteria, allowing internal gaps of up to two residues. Drug abbreviations: CAS, caspofungin; MCF, micafungin; AND, anidulafungin; RZF, rezafungin; IBX, ibrexafungerp; ECN, echinocandin.

Comparing selection coefficient distributions across the targeted regions, we observed that resistance-associated substitutions were not uniformly distributed but concentrated in discrete resistance regions (**Figure 1C**). For each amino acid substitution and condition, we first averaged barcode-level selection coefficients within each independently constructed library and then averaged the library means with equal weight, yielding a two-step average referred to here as the nested mean selection coefficient (*S*). Next, selection coefficients were centered relative to the wild-type amino acid at the same position and scaled within each drug-region combination. We then plotted, for each position, the fraction of substitutions with strong positive selection (*S*_scaled_ > 0.5), upon drug exposure, and the fraction of substitutions with strong negative selection (*S*_scaled_ < −0.5) for the no-drug condition. This representation highlights where multiple amino acid substitutions at the same position repeatedly conferred resistance under drug selection, rather than focusing on individual high-scoring alleles (see below). We defined resistance regions from the DMS data as clusters of adjacent positions at which multiple substitutions had strong positive selection coefficients. For echinocandins, we used a mean smoothed resistance fraction of at least 0.15 across CAS, AND, and RZF, with at least two of these three echinocandins independently crossing a fraction threshold of 0.125. Short internal gaps of up to two residues were allowed, because resistance hotspots are expected to represent clusters rather than perfectly uninterrupted regions of resistance-prone residues. MCF was plotted but was not used for ECN-resistance region calling because its overall signal was weaker, especially in HS3, indicating a lower effective selection under the MCF screening condition.

Using this definition, the DMS-derived ECN-resistance regions overlapped the historical canonical hotspot regions but extended beyond them in several cases. In the HS1-region, the ECN-resistance region spanned A631 to R649, extending N-terminally relative to the historical HS1 interval E635 to R649. In the HS3-region, ECN activity spanned I686 to N700, extending upstream of the historical HS3 interval L690 to N700. In the HS2-region, ECN activity covered D1353 to V1365, including the historical HS2 interval D1353 to S1361 and extending several residues downstream. Thus, the DMS data support the historical hotspot definitions but indicate that the experimentally observed echinocandin resistance landscape is broader than the historical canonical boundaries. Applying the same position-level framework to IBX revealed both overlap and divergence from the ECN-resistance regions. IBX resistance overlapped with the ECN-resistance regions in HS1 and HS3 but was more restricted, spanning A631 to S643 in HS1 and L687 to I698 in HS3. The strongest difference was observed in the HS2-region, where IBX activity extended from S1361 to I1373. Residues S1361 to V1365 were within both ECN- and IBX-resistance regions, but V1366 to I1373 was biased to IBX resistance.

Defining hotspot boundaries determines which *FKS1* positions are included in targeted genotyping and how detected variants are interpreted. Restricting *FKS1* variant analyses to the historical regions could miss adjacent resistance-associated substitutions. We therefore propose to use our DMS-defined resistance regions to refine hotspot terminology while distinguishing echinocandin-associated and IBX-biased effects. For echinocandins, we propose extending the operational boundaries of the established hotspots to the full ECN-resistance regions identified here: HS1 from A631 to R649, HS3 from I686 to N700, and HS2 from D1353 to V1365. These revised intervals retain the historical hotspot boundaries while including adjacent residues with recurrent DMS-supported resistance activity, and they also encompass the overlapping IBX-responsive positions in HS1, HS3, and the proximal HS2-region. However, because IBX activity in the HS2-region extended further downstream and was strongly biased toward the triterpenoid condition, we propose designating I1362 to I1373 as HS4 rather than simply extending HS2. This distinction preserves the clinically established meaning of HS2 for echinocandin resistance while recognizing a partially overlapping, triterpenoid-biased resistance region. Under this nomenclature, HS2 and HS4 overlap at I1362 to V1365, but their relevance differs by drug class.

### Mutation-level profiles separate cross-resistant from drug-specific resistant substitutions

Next, we analyzed selection coefficients for individual mutations per position. Under drug exposure, enrichment is interpreted here as evidence of reduced susceptibility. In **Figure 2**, we plotted the nested mean selection coefficient (*S*) for every single amino acid substitution across the three targeted Fks1 regions for all drugs tested (**Figure 2A**), and compared these mutation-level profiles between IBX and AND, as an ECN representative (**Figure 2B**). For this analysis, we zoomed in on the ECN- and IBX-resistance regions defined in **Figure 1C**, extended by seven residues on both sides to retain immediately adjacent substitutions with potential resistance activity. This captured the full resistance-enriched segments while improving visualization of individual mutation effects. Full heatmaps across the complete DMS-targeted regions and all pairwise echinocandin-IBX comparisons are shown in **Supplementary Figures S6** and **S7**, respectively.

**Figure 2.**
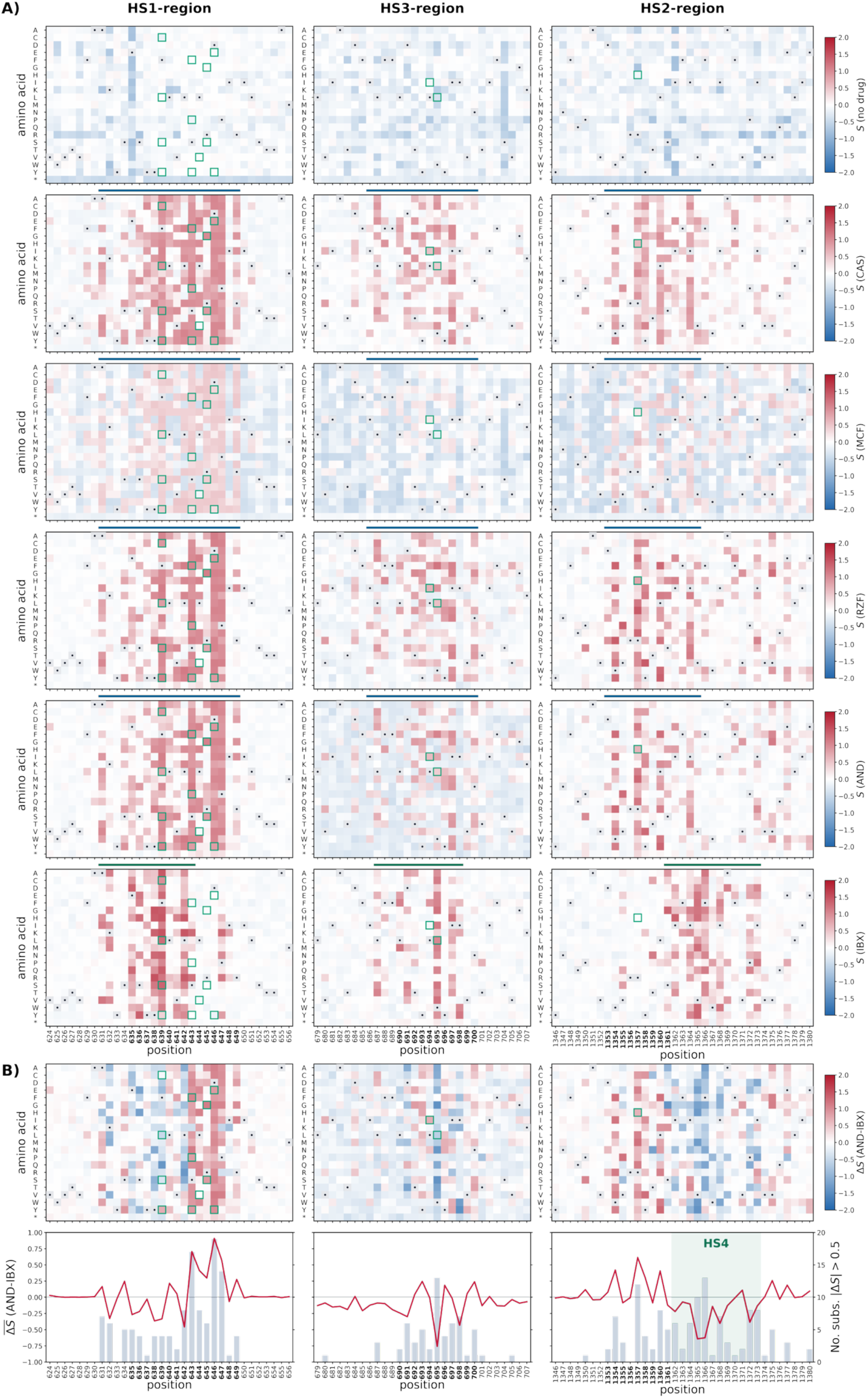
Mutation-level resistance landscapes identify cross- and drug-specific resistance substitutions. **A)** Selection coefficients (*S*) averaged first across synonymous barcode estimates within each independently constructed library and then equally across library means (nested means) for all single amino acid substitutions across the ECN- and IBX-resistance regions identified in **Figure 1C**, extended by seven residues upstream and downstream. Full heatmaps across the complete DMS-targeted regions are shown in **Supplementary Figure S6**. Rows show no drug, CAS, MCF, RZF, AND, and IBX. Red indicates positive selection coefficients and blue indicates negative selection coefficients, calculated from changes relative to the matched input population (t0) and normalized to silent variants. Wild-type residues are shown as black dots on light grey tiles, and priority substitutions identified from FungAMR^16^ and the Gifford *et al. C. auris* dataset^6^ are outlined in green. Dark blue and green bars indicate ECN- and IBX-resistance regions, respectively; historical hotspot positions are indicated by bold black residue numbers. **B)** Differential selection between AND and IBX across the same regions. Heatmaps show Δ*S*_*m,AND−IBX*_ = *S*_*m,AND*_ −*S*_*m,IBX*,_where positive values indicate higher selection coefficients under AND and negative values indicate higher coefficients under IBX. Line plots show the position-wise mean differential selection coefficient .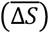 on the left y-axis, and grey bars show the number of substitutions at each position with |Δ*S* | *>* 0.*5*, on the right y-axis. The green shaded area marks HS4. Equivalent echinocandin-versus-IBX comparisons for CAS, MCF, RZF, and AND are shown in **Supplementary Figure S7**.

In addition, we focused our analysis on priority substitutions identified from recurrent fungal pathogen variation and curated resistance annotations, using the *C. auris* population dataset of Gifford *et al*.^6^ and the FungAMR database^16^ (**Supplementary Figure S8** and **Supplementary Table S2**). These substitutions were marked on the heatmaps to directly compare experimentally measured resistance effects with variants observed in fungal pathogens. Overall, the mutation-level heatmaps confirmed the position-level patterns observed in **Figure 1C**, but also resolved which specific substitutions drove cross-resistance versus drug-biased resistance.

In the HS1-region, substitutions at F639 showed the clearest cross-resistance profile. F639C and F639L were enriched under both echinocandins and IBX exposure, consistent with cross-resistance. F639L had *S* = 1.27 under IBX and *S* = 0.78-0.94 across AND, RZF, and CAS. F639C behaved similarly, with *S* = 0.82-0.90 across the main echinocandins and *S* = 0.97 under IBX. In contrast, several substitutions in the central and C-terminal part of HS1 were strongly biased towards echinocandin resistance: S643F, S643P, S643Y, and D646Y consistently had positive selection coefficients under echinocandins but near-zero or negative coefficients under IBX. For example, S643F reached *S* = 0.98-1.05 across CAS, RZF, and AND, but had a near-zero coefficient under IBX (*S* = -0.04). Similarly, D646Y was enriched under echinocandins (*S* = 0.80-0.86) and depleted under IBX (*S* = -0.15). These substitutions therefore explain why the ECN-resistance region spans the full HS1-region, whereas the IBX-resistance region is more restricted (**Figure 1C**). The extended HS1 boundary was also supported by sequence divergence in fungal pathogens. In the Gifford *et al*.^6^ dataset, *C. auris* A627T and A627V mapped to the N-terminal extension of the *S. cerevisiae* HS1-region. The corresponding *S. cerevisiae* substitutions A631T and A631V had positive selection coefficients in the DMS maps. A631V was enriched under both echinocandins and IBX (*S* = 0.95 for CAS, 1.02 for RZF, 1.00 for AND, and 1.16 for IBX), while A631T was positively selected under most echinocandins (*S* = 0.86-0.88 for CAS, RZF, and AND). This supports the inclusion of residues upstream of the historical HS1 boundary in the revised hotspot definition.

In the HS3-region, only two priority substitutions were detected, but the available mutations again separated cross-resistance and drug-biased effects. L694I was enriched under echinocandins, most strongly under RZF (*S* = 0.65), but depleted under IBX (*S* = -0.11). W695L showed the opposite pattern and was the clearest IBX-biased priority substitution in HS3, with a higher selection coefficient under IBX (*S* = 1.02) than under the echinocandins AND, CAS, and RZF (*S* = 0.41 for AND, 0.44 for CAS, and 0.54 for RZF) and a negative coefficient under MCF (*S* = -0.17). Thus, HS3 contains overlapping ECN and IBX resistance potential, but the individual substitutions contributing to each drug class are not identical.

The HS2-region showed the strongest separation between echinocandin and IBX resistance. Substitutions at R1357 were enriched under echinocandins but not under IBX. R1357H showed *S* = 0.61 for CAS, 0.77 for RZF, and 0.63 for AND, but had a near-zero coefficient under IBX (*S* = -0.02). R1357Y followed the same pattern, with weaker enrichment under echinocandins and no enrichment under IBX. Conversely, the substitutions with the highest selection coefficients under IBX were concentrated downstream of HS2, within the HS4 region defined in **Figure 1C**. The AND-IBX differential heatmap and position-level comparison in **Figure 2B** illustrate this shift: positive values indicate higher selection coefficients under AND, whereas negative values indicate higher coefficients under IBX. In HS2, the strongest negative differential signal clustered around the downstream segment corresponding to HS4, supporting a triterpenoid-biased resistance region distinct from the canonical HS2. The extended HS2 boundary was also supported by pathogen variation. The *C. auris* substitution I1361T identified in the Gifford *et al*.^6^ dataset mapped to *S. cerevisiae* I1364T, which was enriched under echinocandins in our DMS data (*S* = 0.84 for CAS, 1.13 for RZF, and 1.00 for AND), but had a near-zero coefficient under IBX (*S* = 0.01). This supports the downstream extension of the ECN-resistance region beyond the historical HS2 boundary. At the same time, individual substitutions outside the updated resistance-region definitions were also observed. For example, P1350S, upstream of HS2, was enriched under echinocandins (*S* = 0.62 for CAS, 1.02 for RZF, and 1.00 for AND), whereas E1377P, downstream of HS4, was enriched under IBX (*S* = 0.75). However, these isolated substitutions were not used to extend hotspot boundaries, because the hotspot definition was based on clusters with recurrent positive selection across multiple substitutions and, for ECNs, across multiple drugs. Their rarity in clinical and population datasets further suggests that such isolated positions are less central to the resistance landscape than the clustered ECN and IBX regions.

The no-drug control showed near-zero selection coefficients, and thus low fitness costs, for most priority substitutions. Among the priority substitutions, no-drug scores generally ranged from *S* = -0.16 to 0.04. The main exception was W695L, which was more depleted in the absence of drug (*S* = -0.37), suggesting a basal fitness cost that is overcome under IBX selection.

### Individual mutant validation confirms differential resistance patterns of selected alleles

To validate the DMS landscapes, we tested selected alleles directly by measuring drug susceptibility in individual *S. cerevisiae* and *C. auris* mutants. We focused mainly on HS1 because this region contains the highest density of clinically relevant echinocandin-resistance substitutions (**Supplementary Figure S8** and **Table S2**) and showed a clear position-specific difference between echinocandin and IBX resistance in the DMS heatmaps (**Figure 2**). We validated selected alleles in 10 HS1 single-amino-acid mutants from Durand *et al*. in *S. cerevisiae*^8^, and in *C. auris* using 13 experimentally evolved mutants, 11 CRISPR-Cas9-engineered mutants, and 19 clinical isolates with Fks1 variation from different geographical origins to account for strain-background variation. Strain information is provided in **Supplementary Table S3**. We tested AND as the representative echinocandin, paired with IBX as in **Figure 2B**. We evaluated drug susceptibility of individual mutants using standardized EUCAST broth dilution assays and inferred changes in susceptibility from the area under the curve (AUC) of the dose–response curve, since no clinical breakpoint MIC values have been described for IBX. For engineered and experimentally evolved strains with matched parental (WT) controls, we quantified relative resistance as the AUC/AUC_max_ of the mutant normalized to its parental wild-type strain. Values above 1 therefore indicate a relative decrease in susceptibility. For clinical isolates without an isogenic parental strain, we used absolute AUC/AUC_max_ values. We used an operational IBX-resistance threshold corresponding to an MIC of 1 µg/mL, informed by the DMS-derived resistance patterns and the relative-resistance validation data. For AND, we used 0.25 µg/mL as the operational resistance threshold, based on the EUCAST clinical breakpoint for resistance to this drug^17^.

The *S. cerevisiae* validation set in **Figure 3A** largely recapitulated the mutation-level DMS patterns. Substitutions at the N-terminal side of HS1, including ScFks1 F639C, produced cross-resistance to AND and IBX, consistent with the shared F639 resistance signal in the DMS landscape. Other validated HS1 substitutions showed drug-specific mutational effects on resistance. ScFks1 L642G increased IBX resistance without a corresponding AND shift, whereas ScFks1 S643F, ScFks1 R645S, and ScFks1 D646W showed AND resistance with only weak or threshold-adjacent IBX effects. These results are consistent with **Figure 2**, where substitutions in the central HS1 segment, including S643 and D646 variants, were enriched under echinocandins, with near-zero or negative selection coefficients under IBX.

**Figure 3.**
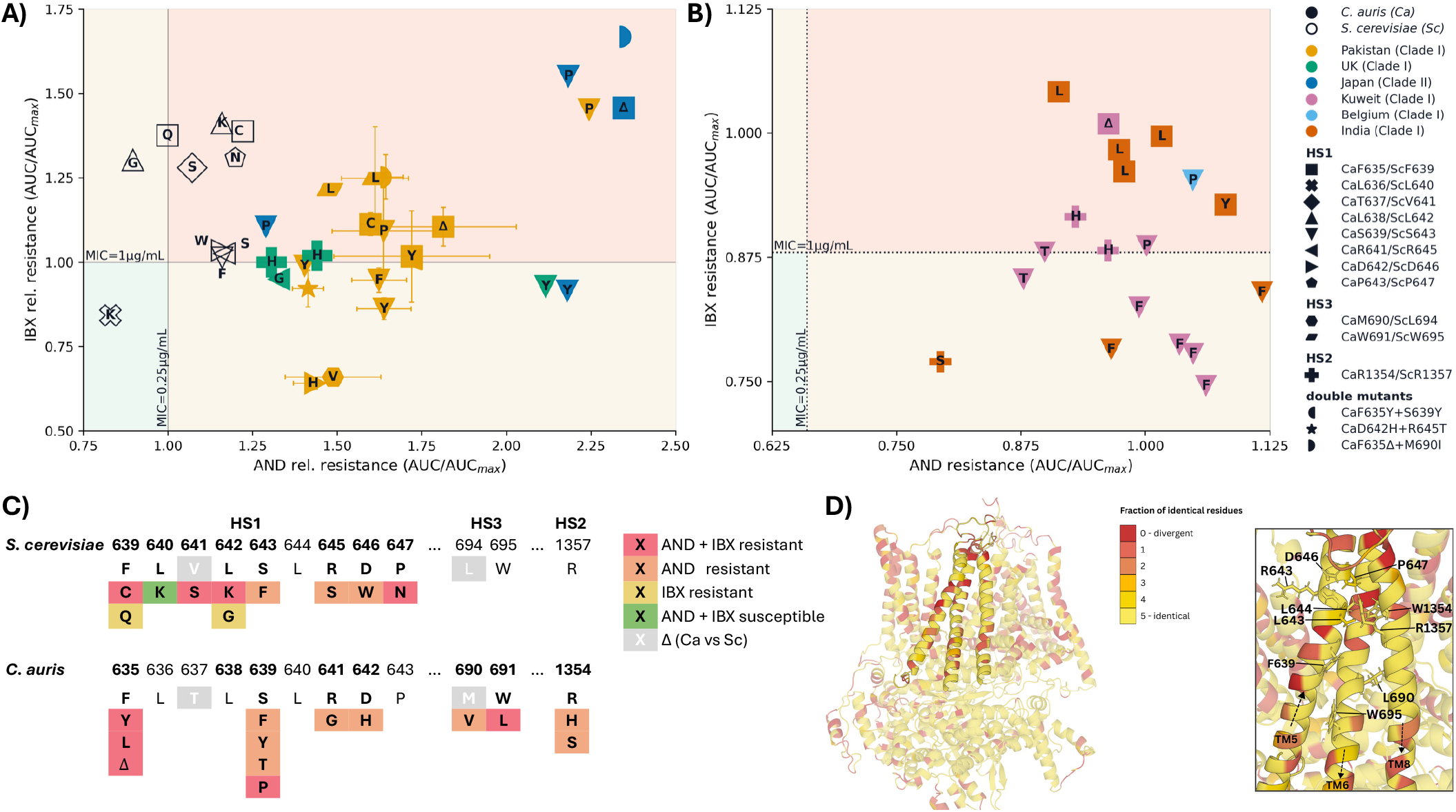
Individual mutant validation of DMS-derived echinocandin and ibrexafungerp resistance patterns. **A**) Relative susceptibility of selected *S. cerevisiae* and *C. auris FKS1* mutants to anidulafungin (AND) and ibrexafungerp (IBX). Values represent AUC/AUC_max_ relative to the corresponding parental wild-type strain. Values above 1 indicate reduced susceptibility. Within the figure, Ca and Sc denote *C. auris* and *S. cerevisiae*, respectively. Symbol shape indicates the Fks1 residue, and the letter inside each symbol indicates the amino-acid substitution, with the exception of double mutants, which are indicated by a unique symbol. Error bars, when shown, indicate the standard error of the mean (SEM) susceptibility value determined by three independently constructed mutants. Symbol colors indicate geographical origin and clade. **B**) Absolute susceptibility of clinical *C. auris FKS1* mutant isolates from Kuwait, Belgium, and India. Because matched parental strains were not available, resistance was inferred from operational MIC-linked AUC/AUC_max_ thresholds: AND = 0.25 µg/mL and IBX = 1 µg/mL. Symbol colors indicate geographical origin and clade. **C**) Summary of validated resistance patterns mapped onto aligned *S. cerevisiae* and *C. auris* Fks1 residues. Colours indicate AND resistance, IBX resistance, cross-resistance, or susceptibility based on the integration of validation data and DMS data. Grey residues indicate amino-acid differences between the two species. **D)** Fks1 sequence conservation between *S. cerevisiae* and five major candidiasis-causing fungal pathogens listed as critical or high priority by the WHO fungal priority pathogens list^18^: *C. auris, C. albicans, N. glabratus, C. parapsilosis*, and *C. tropicalis*. Residues are colored on the *S. cerevisiae* Fks1 structure according to the number of pathogen orthologues with an identical amino acid at the aligned position. Residues highlighted on the Fks1 structure indicate the 10 most frequently reported echinocandin-resistance-associated positions in FungAMR^16^, mapped to the corresponding *S. cerevisiae* Fks1 residues, summarized in **Supplementary Table S1**.

The *C. auris* validation data in **Figure 3A-B** showed the same broad structure, but with minor allele- and background-specific differences. CauFks1 F635 substitutions were generally cross-resistant, matching the shared resistance behavior of the corresponding ScFks1 F639 position. CauFks1 S639F, S639Y, and S639T mutants showed AND resistance without strong IBX resistance, consistent with the echinocandin-biased DMS signal at the corresponding *S. cerevisiae* position. However, CauFks1 S639P was an exception: CRISPR-engineered, experimentally evolved, and clinical CauFks1 S639P mutants all fell in the cross-resistant region, despite the corresponding ScFks1 S643P allele being echinocandin-biased in the DMS map.

Additional validated alleles beyond HS1, were in agreement with DMS data. CauFks1 M690V in HS3 was AND-resistant without strong IBX resistance, whereas CauFks1 W691L was cross-resistant, consistent with the DMS prediction that HS3 contains overlapping but substitution-specific resistance determinants. CauFks1 R1354H and R1354S, corresponding to the echinocandin-associated HS2 residue ScFks1 R1357, were predominantly AND-resistant, although some measurements were close to the operational IBX threshold.

Together, these validations show that the DMS maps capture resistance effects that are reproducible in individual mutants and informative across species (**Figure 3C**). Thus, despite specific exceptions such as CauFks1 S639P, *S. cerevisiae* provides a powerful model for defining mutational resistance landscapes in conserved fungal drug targets such as Fks1. To assess how broadly this model may extend across clinically important fungi, we next mapped Fks1 sequence conservation between *S. cerevisiae* and five major candidiasis-causing pathogens from the critical and high-priority groups of the WHO fungal priority pathogens list^18^: *C. auris, Candida albicans, Nakaseomyces glabratus, Candida parapsilosis*, and *Candida tropicalis* (**Figure 3D**). Full-length Fks1 identity ranged from 72.49% to 72.87% between *S. cerevisiae* and the four *Candida/Candidozyma* species, and reached 86.30% for *N. glabratus*. Across all five pathogens, 64.71% of aligned residues were identical to *S. cerevisiae*. This conservation was not uniform across the protein, but the resistance-relevant regions were generally well conserved. Most validated positions in HS1, HS3, and HS2 were identical between *S. cerevisiae* and *C. auris*, including the major resistance-associated positions ScF639/CauF635, ScS643/CauS639, ScR645/CauR641, ScD646/CauD642, ScW695/CauW691, and ScR1357/CauR1354 (**Figure 3C-D**). The main differences occurred at a subset of neighboring positions, such as ScV641/CauT637 and ScL694/CauM690, emphasizing that conservation is high but not complete. This pattern explains why the *S. cerevisiae* DMS landscape predicts most pathogen-relevant resistance outcomes, while also allowing species- or allele-specific deviations such as the cross-resistance observed for CauFks1 S639P.

### Distinct interaction networks provide a structural basis for differential resistance patterns

To investigate the molecular basis of the differential resistance landscapes, we generated structural models of the Fks1-glucan complex with each of the five inhibitors. Initial ligand poses were obtained by structure-guided placement and refined by MD-based simulated annealing using the GBSW implicit-solvent and membrane model. Candidate poses were selected based on their consistency with DMS-defined resistance patterns, structural agreement with the cryo-EM Fks1-glucan-CAS complex (PDB 9PE1)^7^, and energetic criteria (**Figure 4A**).

**Figure 4.**
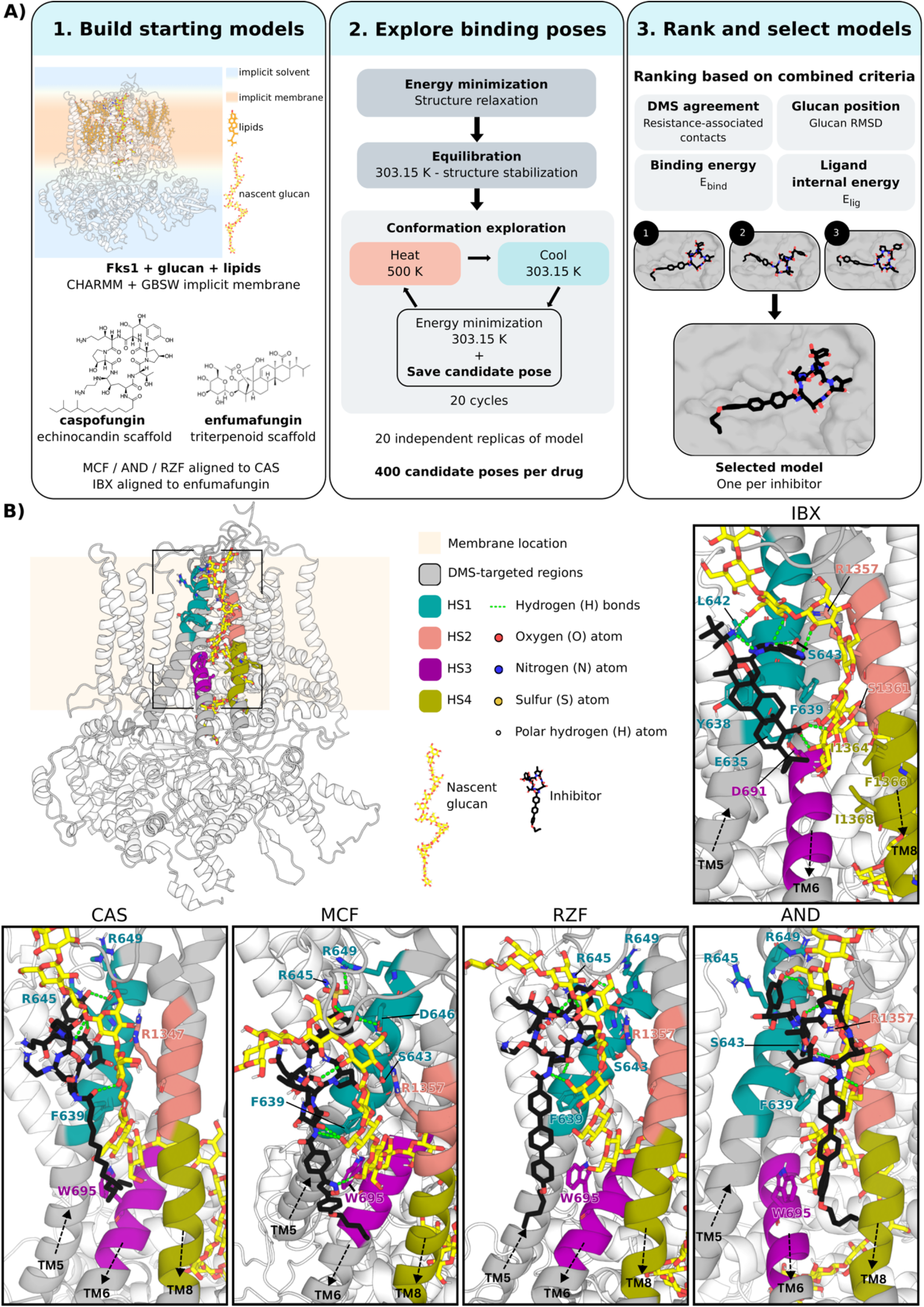
Predicted binding modes of echinocandins and IBX in the Fks1-glucan complex. **A)** Overview of the MD-based simulated-annealing and pose-selection workflow. Structural models of Fks1–glucan complexes were constructed from the 3.09 Å cryo-EM structure of the CAS-bound complex (PDB 9PE1)^7^. Missing loops were modeled, and the complexes were built with CHARMM. The experimentally determined CAS pose^7^ was retained as the initial reference, while the initial positions of the other echinocandins were obtained by structural superposition onto CAS. The initial IBX pose was generated by superposition onto enfumafungin from its cryo-EM complex (PDB 9WZU)^11^. Following energy minimization and equilibration, MD-based simulated annealing was performed through repeated heating (500 K) and cooling (303.15 K) cycles to enhance conformational sampling. Twenty independent replicates of 20 annealing cycles each generated 400 candidate poses per inhibitor, with one pose retained after each cycle. Candidate poses were ranked using combined structural, experimental, and energetic criteria: *DMS_residues*, the number of DMS-identified resistance-associated residues in contact with the drug; *Glucan_RMSD*, the root-mean-square deviation of glucan after protein alignment, used to assess preservation of its position within the translocation channel; *E*_*lig*_, the internal energy of the ligand; *E*_*bind*_, the ligand binding energy calculated as the energy of the Fks1-glucan-ligand complex minus the energies of the isolated Fks1-glucan complex and ligand. **B)** Top-ranked predicted binding modes of IBX, CAS, MCF, RZF and AND. Close-up views illustrate the selected binding pose of each inhibitor within the Fks1–glucan complex. DMS-identified resistance-associated residues located near the ligand–glucan complex are shown as sticks and labeled, with colors indicating their respective hotspot regions. The four echinocandins adopt similar predicted binding geometries, with their cyclic hexapeptide cores interacting with Fks1 and the nascent glucan and their hydrophobic acyl side chains extending toward the membrane. In contrast, IBX adopts a more compact binding mode with a distinct interaction network. In all five models, the inhibitors are positioned near the glucan extrusion site and form predicted contacts with both Fks1 and nascent glucan, which remains within the translocation channel. Two-dimensional ligand interaction diagrams are provided in **Supplementary Figure S9**.

The four echinocandins adopted similar predicted binding modes at the extracellular membrane-protein interface (**Figure 4B**), consistent with their shared cyclic hexapeptide structure. The macrocycle and the proximal acyl side chain were positioned alongside the nascent glucan, which occupies much of the groove formed by the Fks1 hotspot regions, while the hydrophobic tail extended toward the membrane. Notably, the simulated-annealing-refined CAS model reproduced key features of the CAS-bound cryo-EM binding mode (PDB 9PE1)^7^, in which the macrocycle engages both Fks1 and glucan, while the lipid tail anchors the drug at the extracellular membrane leaflet. The four refined models shared a network of predicted interactions involving hydrogen bonds between the homotyrosine hydroxyl group and residues D646 and K1336 (through its backbone in the MCF model), a cation–π interaction between the homotyrosine aromatic ring and R645, and hydrogen bonds involving S643 and glucan. Additional hydrogen bonds between the acyl linker and glucan hydroxyl groups varied with ligand orientation.

Despite this conserved binding mode, the echinocandins exhibited compound-specific differences. The aliphatic acyl chain of CAS extended less deeply into the membrane-facing groove than the bulkier aromatic side chains of the other echinocandins. Its ethylenediamine and 3-hydroxyornithine substituents faced the solvent-membrane interface without forming direct interactions with Fks1 or glucan, although the intramolecular hydrogen bond involving the 3-hydroxyornithine amino group contributed to the predicted ligand conformation. AND and RZF adopted closely related binding geometries, consistent with their shared acyl side chain and the substitution of the ornithine hemiaminal hydroxyl group in AND by a choline aminal ether in RZF. Their differences were largely restricted to the orientation of this substituent and minor variations in acyl chain positioning. In contrast, MCF exhibited a distinct polar interaction network: its homotyrosine sulfate group interacted with R645 and R649, while heteroatoms of the isoxazole ring in its acyl substituent formed predicted hydrogen bonds with glucan.

IBX adopted a more compact binding geometry, characterized by fewer direct contacts with Fks1 and involving a distinct network of interactions with nascent glucan (**Figure 4B**). Its rigid polycyclic triterpenoid scaffold occupied a membrane-facing region near a sterol-associated site observed in PDB 9PE1^7^. Notably, the carboxyl group formed two predicted hydrogen bonds with two glucan hydroxyl groups deep within the membrane-facing groove, at a depth comparable to the isoxazole ring of MCF. Additional hydrogen bonds involved the pyridine nitrogen, the 1,2,4-triazole ring, and the terminal aminoalkoxy group, primarily with glucan. One triazole nitrogen also interacted with S643, while another formed an intramolecular hydrogen bond with the terminal amino group of the aminoalkoxy substituent, potentially stabilizing the ligand conformation. Most direct IBX-Fks1 contacts were concentrated near the N-terminal portion of HS1. This localization is broadly consistent with cryo-EM studies of enfumafungin, which identify TM5 as a major component of its binding environment^11^. However, the extensive IBX–glucan interactions predicted here suggest an additional role for nascent glucan in stabilizing the inhibitor-bound complex, a possibility that remains to be experimentally established.

Together, these predicted binding modes provide a structural framework for interpreting the shared and drug-specific resistance patterns revealed by DMS. Three residues associated with resistance to all five inhibitors (A631, F639 and W695) may contribute through distinct mechanisms. A631 lies behind W695, which lines the glucan translocation groove, suggesting that substitutions at A631 could indirectly influence inhibitor binding by altering W695 positioning. In contrast, F639 and W695 are positioned near both glucan and the inhibitors, where substitutions could more directly affect the local ligand binding environment. Five additional residues (S643, D646, L694, W1354 and R1357) were associated with resistance to all four echinocandins but not IBX, consistent with the more extensive Fks1-glucan-macrocycle interaction network predicted for echinocandins. S643 and D646 form direct hydrogen bonds with the echinocandin macrocycle, while R1357 is positioned to influence the organization of the glucan-binding interface. L694 contributes to the glucan-occupied groove, whereas W1354, buried behind R1357 within HS2, may exert more indirect structural effects. Although IBX also forms a predicted hydrogen bond with S643, the predominantly echinocandin-associated DMS phenotype at this position suggests that similar contacts do not necessarily confer equivalent contributions to drug susceptibility. These observations suggest that resistance may arise through both disruption of direct drug interactions and changes in the Fks1-glucan architecture required for inhibitor binding.

## Discussion

Our study identifies three main findings: (1) resistance-associated variation extends beyond the canonical *FKS* hotspot boundaries, motivating revised intervals for surveillance; (2) echinocandins and ibrexafungerp exhibit overlapping yet asymmetric mutational resistance landscapes and validation of selected alleles in *C. auris* supports the majority of the drug-specific patterns identified in *S. cerevisiae*; and (3) structural models incorporating nascent glucan suggest that distinct inhibitor–protein–glucan interactions could explain these asymmetries.

We propose to extend the hotspot boundaries because the DMS-derived resistance enrichment regions did not coincide with the canonical intervals. Echinocandin-associated enrichment extended upstream to A631 in the HS1 region, upstream to I686 in the HS3 region, and downstream to V1365 in the HS2 region, whereas IBX-associated enrichment in the latter region shifted further downstream, from S1361 to I1373, which led us to designate I1362 to I1373 as HS4 rather than simply widening HS2. Because these newly assigned intervals were derived from one species under laboratory selection, their relevance to clinical surveillance requires evaluation in pathogenic fungi. However, published datasets provide examples of variation in these extended regions in pathogens. The survey of 12,644 *C. auris* genomes by Gifford *et al*. identified five substitutions lying outside the hotspots but within our proposed boundaries: A627T and A627V (ScFks1 A631), I1361T (ScFks1 I1364), and F1367C and F1367L (ScFks1 F1370)^6^, although their contribution to reduced susceptibility was not established in that study. In addition, the extension of HS1 is supported by the experimental evolution study of Ksiezopolska *et al*., in which three echinocandin-resistant clones of *N. glabratus* were recovered with Fks2 A651T (which maps to ScFks1 A631T) upstream of HS1, with whole genome sequencing excluding other candidate responsible loci for acquired echinocandin resistance^19^. For the HS3 extension, Fernández-Silva *et al*. described a previously unreported Fks2 L707S (which maps to ScFks1 L687S) in six *N. glabratus* strains^20^, some of which had an increased AND or CAS MIC. Downstream of the traditional HS2 and in HS4, the amino acid corresponding to ScFks1 I1364 was found mutated in at least four species: in a *C. auris* clinical isolate, I1361T was tentatively associated with CAS resistance^21^, *C. albicans* I1368T was one of only two out-of-hotspot substitutions among 192 echinocandin-resistant isolates^12^, an echinocandin-resistant *C. tropicalis* isolate carried I1368S together with F641L^22^, and an isolate of *Diutina catenulata* (formerly *Candida catenulata*) with Fks1 I1348S was recovered with echinocandin MICs of 4 µg/mL or higher^23^. Ren *et al*. report CAS resistance for ScFks1 V1365F, downstream of the historical HS2 region^7^. This is one of the few mutations scored as resistant to both classes in our DMS screen. These observations are consistent with the proposed downstream extension of HS2, although the independent effects of these substitutions remain incompletely resolved. Clinical variation also occurs at the position corresponding to ScFks1 F1370, where the variants discussed here were not enriched in our DMS screens: beyond the two CauFks1 F1367 variants reported by Gifford *et al*.^6^, *C. parapsilosis* Fks1 F1386S^24^ and *D. catenulata* Fks1 F1354L^25^ were identified in clinical isolates. The effect of these mutations on susceptibility was not individually validated by genetic engineering, except for *C. parapsilosis* Fks1 F1386S, revealing an interesting insight: introducing F1386S into a susceptible *C. parapsilosis* background by CRISPR-Cas9 left all three echinocandin MICs at or below those of the parental strain while conferring high MCF tolerance, based on survival in 4 µg/mL of MCF^26^. A Brazilian *C. parapsilosis* candidemia outbreak provided further functional evidence for non-hotspot *FKS1* variation (Fks1 E1393G) affecting tolerance without increasing resistance^27^. Since our pooled growth assays select for increased growth at otherwise inhibitory concentrations and thus resistance, the effect of increased survival and thus tolerance of a mutant is unknown. This could explain why the corresponding mutant was not enriched in our DMS approach. Our results and previous reports combined support extending the hotspot boundaries in *FKS* genes and broadening sequence coverage for epidemiological surveillance. Nevertheless, resistance-associated variation can even occur beyond these revised intervals, although it often concerns individual substitutions rather than positions at which multiple substitutions confer resistance. For example, an *N. glabratus* isolate carrying Fks2 P1371S was resistant to CAS but susceptible to MCF and AND, and an echinocandin-resistant *C. albicans* isolate carried Fks1 P1354S^28^. Both substitutions correspond to ScFks1 P1350S, which was enriched under echinocandins in our screens. However, enrichment across substitutions at this position did not meet our criteria for inclusion within the revised hotspot boundaries.

Our results identify conserved, position-dependent differences between echinocandin and triterpenoid susceptibility. Comparisons of literature reports with our DMS enrichment and individual-mutant validation require two distinctions. MIC differences between unrelated clinical isolates do not separate the contribution of an *FKS* mutation from strain-background effects, and clinical breakpoints have not been established for IBX, although wild-type upper limits based on MIC distributions have been proposed for some species (e.g. *C. albicans* 0.25 µg/mL and *N. glabratus* 0.5 µg/mL, none reported for *C. auris*)^29^. Nevertheless, several literature reports align with our observations of cross-resistance or drug-biased patterns. At ScFks1 F639, our cross-class enrichment and *C. auris* F635 validation agree with parent–descendant comparisons in equivalent *N. glabratus* Fks1 mutants F625L and F625Δ that showed increased IBX and MCF MICs^13^. Jacobs *et al*. tested 12 clinical *C. auris* F635C mutants, which all showed AND MICs of 4 µg/mL and high IBX MICs in most isolates: 9 isolates had an MIC of 2 µg/mL, two an MIC of 1 µg/mL and one an MIC of 0.25 µg/mL^30^. However, de Groot *et al*. and Jørgensen *et al*. classified a *C. auris* Fks1 F635Y isolate as echinocandin resistant but not IBX resistant, with IBX MICs of 0.5 µg/mL and 1 µg/mL, respectively^29,31^. Likewise, Díaz-García *et al*. reported an echinocandin-resistant *C. albicans* F641L (equivalent to ScFks1 F639L) isolate with IBX MIC 0.125 µg/mL^32^. Thus, this conserved position supports cross-class susceptibility shifts without uniformly producing an elevated IBX MIC in all species, strains and studies.

At ScFks1 S643, our *C. auris* S639F/Y validation aligns closely with Arendrup *et al*. and Jacobs *et al*.: their isolates had elevated echinocandin MICs but low IBX MICs of 0.25–0.5 µg/mL^30,33^. S639P was an important exception in our validation, reducing susceptibility to both AND and IBX despite the echinocandin-biased ScFks1 S643P DMS phenotype. Likewise, Spruijtenburg *et al*. classified three out of four *C. auris* S639P variants as resistant to both SCY-247 (a second-generation triterpenoid analogue of IBX) and MCF, while one was resistant only to MCF^34^. In the same study, four S639F isolates were classified as resistant to MCF but not SCY-247, aligning with our results, although one S639T mutant and one S639Y mutant were classified as cross-resistant, conflicting with our findings. In addition, You *et al*. measured an approximately 15-fold increase in enfumafungin MIC for S643P in *S. cerevisiae*^11^. The IBX susceptibility phenotype of mutations at ScFks1 S643/CauFks1 S639 therefore cannot be extended to all triterpenoids, and species- or strain-specific differences in susceptibility for the same mutation exist.

Further along HS1, clinical evidence largely supports the echinocandin-biased DMS profiles at ScFks1 R645 and D646. Nunnally *et al*. reported low (<0.03 – 0.5 µg/mL) IBX MICs for *N. glabratus* Fks1 R631G and D632V/Y (which map to ScFks1 R645 and D646, respectively) mutants^35^, while Díaz-García *et al*. explicitly classified *N. glabratus* Fks2 D666E/G (which map to ScFks1 D646) isolates as echinocandin resistant but IBX wild type^36^. Nevertheless, Pfaller *et al*. reported an IBX MIC of 4 µg/mL for an echinocandin-resistant *N. glabratus* isolate with Fks1 D632Y^37^.

For ScFks1 W695 in HS3, Martins *et al*. provided direct allele-level evidence: reconstructing W760S in Bgs4 (which maps to ScFks1 W695), the *Schizosaccharomyces pombe* orthologue of Fks1, reduced susceptibility to enfumafungin and CAS^38^, agreeing with the drug-dependent ScFks1 W695S profile of our DMS results. Similarly, *N. glabratus* Fks2 W715L (which maps to ScFks1 W695) increased IBX and MCF MICs approximately eight- and tenfold relative to its parent^13^. ScFks1 W695L conferred resistance to IBX, CAS, AND and RZF in our DMS analysis. At ScFks1 R1357, the predominantly AND-associated effects of our *C. auris* R1354H/S mutants and echinocandin-biased DMS profiles agree with low IBX MICs reported for *C. albicans* R1361H/S^29,32^. Jiménez-Ortigosa *et al*. previously identified *N. glabratus* Fks2 A1390D, equivalent to ScFks1 A1369D in HS4: IBX and MCF MICs increased 8- and 1.5-fold, respectively, while enzyme IC_50_ increased approximately 20-fold for IBX and remained unchanged for MCF^13^. In our DMS analysis, A1369D is also IBX-biased, with *S* = 1.07 under IBX compared with *S* = 0.15–0.70 under echinocandin exposure.

Overall, the majority (16 of 22) of unambiguous mutation-level literature comparisons discussed here were directionally concordant with our DMS and individual-mutant validation, while three additional comparisons yielded mixed or non-comparable evidence.

To investigate the molecular basis of differential resistance, we combined structure-guided modeling, MD-based simulated annealing, and DMS-guided pose selection. Our glucan-containing models suggest that differences between echinocandin and triterpenoid recognition extend beyond their contacts with Fks1 to interactions with the nascent glucan polymer. Ren *et al*. provide the experimental foundation for this concept by resolving a ternary Fks1-glucan-CAS complex in which the echinocandin macrocycle interacts with both Fks1 and the emerging glucan, and its lipid substituent extends towards the membrane^7^. Their MD simulations further showed that CAS reduces the conformational flexibility of the extracellular glucan segment, supporting a proposed mechanism in which the inhibitor restricts polymer translocation. Thus, nascent glucan constitutes an integral component of the CAS-binding interface rather than simply an adjacent reaction product^7^. Using this experimentally determined complex as a template, our models extend the proposed binding architecture to MCF, AND and RZF. All four echinocandins share predicted interaction networks indicating a common echinocandin-binding architecture, whereas differences in peripheral substituents may account for compound-specific interactions, consistent with the observations of Durand *et al*.^8^. The predicted poses of MCF, AND and RZF, however, remain to be experimentally validated.

The structurally distinct triterpenoid IBX adopted a more compact binding geometry, which can be compared with recent cryo-EM structures of Fks1 bound to its parent compound, enfumafungin. You *et al*. showed that enfumafungin associates primarily with the transmembrane helix TM5 surface of *S. cerevisiae* Fks1 and proposed that its binding reorganizes the surrounding lipid environment and stabilizes a basal conformation^11^. Similarly, Yang *et al*. reported a predominantly hydrophobic binding interface involving TM5–TM6 in *N. glabratus* Fks1^10^. Unlike the CAS-bound structure described by Ren et al.,^7^ neither enfumafungin-bound reconstruction resolved a nascent glucan polymer. Although You *et al*. illustrated a composite model incorporating glucan from an earlier Fks1 structure^11^, Ren *et al*. subsequently showed that the previously proposed glucan position was incompatible with key structural elements of their catalytically relevant Fks1 conformations^7^. Our approach examines IBX binding within the experimentally resolved, glucan-engaged Fks1 architecture of Ren *et al*.^7^, allowing us to explore potential interactions absent from the available enfumafungin-bound structures.

The predicted binding geometries provide a molecular rationale for several DMS-defined resistance patterns. Compared with IBX, the larger echinocandin macrocycles form more extensive contacts with the C-terminal region of HS1, particularly L644, R645 and D646, consistent with echinocandin-biased phenotypes associated with substitutions at these positions. S643 also exhibits an echinocandin-biased phenotype despite forming predicted interactions with both drug classes, illustrating that similar molecular contacts do not necessarily contribute equally to drug susceptibility. In contrast, F639 occupies a central position in the binding environment of both echinocandins and IBX, consistent with the prominent cross-resistance observed at this position in our DMS and validation analysis. This interpretation agrees with the stabilizing role of F639 in enfumafungin recognition and the resistance phenotype reported for F639S/A by You *et al*. in *S. cerevisiae*^11^, as well as the involvement of the equivalent F625 residue in *N. glabratus* reported by Yang *et al*.^10^. Ren *et al*. also implicated F639 in CAS binding and report resistance associated with F639I/S^7^. Similarly, W695, within HS3 near the predicted IBX-binding region, was associated with IBX resistance in our DMS and validation experiments, consistent with the enfumafungin results reported by You *et al*.^11^. Both F639 and W695 contribute to the architecture of the glucan translocation channel. Their substitution could therefore influence inhibitor susceptibility through changes in the local binding environment, glucan positioning, or both, rather than exclusively through disruption of direct drug contacts.

A distinctive feature of our IBX model is the extensive predicted interaction network with nascent glucan. Whereas the echinocandin macrocycles contact both Fks1 and glucan, the more compact IBX scaffold makes fewer direct protein contacts outside the N-terminal region of HS1. Instead, several of its polar groups interact with the modeled glucan chain, including its carboxyl oxygens, pyridine nitrogen, triazole nitrogen, and aminoalkoxy group. Particularly notable are two predicted hydrogen bonds between the IBX carboxyl group and glucan hydroxyl groups deep within the membrane-facing groove. These interactions could stabilize the polar carboxyl group in an otherwise unfavorable hydrophobic environment while contributing to IBX recognition. The glucan chain also occupies much of the cavity surrounding the predicted IBX-binding site, suggesting that its positioning may be critical for accommodating the inhibitor. This possibility offers an interpretation of the triterpenoid-biased resistance associated with HS4, whose residues face the glucan pathway but do not directly contact IBX in our selected model. Mutations in this region could alter glucan positioning and indirectly disrupt IBX binding without contacting the inhibitor itself. This proposed glucan-mediated contribution differs from, but is not necessarily incompatible with, the lipid-mediated conformational mechanism suggested for enfumafungin^10,11.^ Together, our DMS results and structural models suggest that antifungal resistance can arise through both direct disruption of inhibitor contacts and indirect changes in the Fks1–glucan architecture. The balance between these contributions may differ between echinocandins and IBX, providing a molecular rationale for their overlapping yet asymmetric resistance landscapes. These asymmetries offer potential therapeutic opportunities, as reduced susceptibility to one inhibitor may leave others active, providing a rationale for resistance-informed antifungal selection and the design of compounds with complementary activity profiles.

## Materials and Methods

### Strains, plasmids, oligonucleotides, culture conditions and drug preparations for DMS

All *Saccharomyces cerevisiae* strains used for DMS were derived from strain R1158 (*URA3*::CMV-tTA *his3Δ1 leu2Δ0 met15Δ0*)^39^. Three recipients carried an hphNT1 (HYG) cassette in place of the 53-residue *FKS1* segment targeted by the HS1, HS2/TM8, or HS3 library. Yeast was grown on YPD agar at 30 °C for 48 h and cultured in synthetic complete (SC) medium overnight (12-14 h) before use. YPD plates containing G418 (200 µg/mL) were used to select pCAS-containing transformants; hygromycin B (250 µg/mL) was used during construction and verification of the HYG-marked recipient strains.

The pCAS-HPH plasmid^40^ encoded Cas9, a yeast G418-resistance marker, and a guide RNA targeting HYG, as listed in **Supplementary Table S4**. It was propagated in *Escherichia coli* MC1061 in lysogeny broth (LB) containing kanamycin. Synthetic donor libraries were ordered as oligo pools (oPools) from Integrated DNA Technologies (IDT). Primer names and sequences, guide RNAs, and plasmids are listed in **Supplementary Table S4**; strain identifiers are listed in **Supplementary Table S3**.

AND, CAS, MCF, RZF, and IBX were prepared as 2 mg/mL stocks in 100% dimethyl sulfoxide (DMSO). For screening, 200 µg/mL intermediate stocks were freshly prepared in DMSO and diluted to 20 µg/mL working stocks in water with 10% DMSO.

### Construction of the *FKS1* library-recipient strain for DMS screens

Recipient strains were constructed by replacing each targeted 53-residue *FKS1* interval with a HYG cassette amplified from pFA-hphNT1 using primers containing locus-specific homology, as listed in **Supplementary Table S4**. Transformants were generated by PEG/LiOAc-mediated integration^41^ and selected on hygromycin B. Correct 5′ and 3′ junctions were tested by colony PCR with one primer outside the recombination interval and one primer within HYG, as listed in **Supplementary Table S4**. The three HYG-marked recipients were subsequently used as substrates for pCAS-HPH-mediated reintegration of the wild-type or mutagenized *FKS1* segment.

### Design of saturation-mutagenesis libraries

Three saturation-mutagenesis designs covered 53 amino-acid positions each (159 positions total): one centered on HS1, one centered on HS3, and one spanning HS2. At each position, all 32 NNK codons were represented. Each designed target codon was coupled to one of four synonymous barcode codons upstream or downstream of the mutagenized sequence, providing four nucleotide-level observations for robustness assessment without altering the encoded protein outside the targeted substitution. Thus, each regional pool contained 53 × 32 × 4 = 6,784 designed codon-barcode sequences, and the three-region design contained 20,352 unique sequences.

The complete three-region design was constructed twice independently. For each construction replicate, the three regional oPools were amplified, transformed, harvested, frozen, and screened separately. The two pools for a given region were not combined before selection, allowing construction-specific effects to be measured independently from barcode-level variation.

### Amplification and CRISPR-Cas9 integration of pooled libraries for DMS screens

IDT oPools were resuspended at 50 µM and serially diluted to 0.05 µM for amplification. Each 25-µL KAPA HiFi reaction contained 0.75 µL oPool, 5 µL 5× KAPA Fidelity buffer, 0.75 µL 10 mM dNTPs, 0.75 µL each 10 µM primer, 0.5 µL polymerase, and 16.5 µL nuclease-free water. Cycling was 98 °C for 3 min; 15 cycles of 98 °C for 30 s, 60 °C for 15 s, and 72 °C for 10 s; 72 °C for 20 s; and a 10 °C hold. Forty-eight PCR products were pooled and purified with AxyPrep MAG PCR Clean-up SPRI beads.

Large-scale CRISPR-Cas9 transformations were performed according to Ryan *et al*.^40^ with minor modifications. 15 mL Falcon tubes contained 400 µL purified donor dDNA pool, 150 µL boiled salmon-sperm carrier DNA (10 mg/mL), 93.75 µL pCAS-HPH (160 ng/µL), 1.35 mL competent cells, and 13.5 mL freshly prepared PLATE solution (40% PEG 3350, 100 mM LiOAc, 10 mM Tris-Cl, and 1 mM EDTA). Reactions were incubated for 35 min at 30 °C and 18 min at 42 °C, centrifuged at 1,845 × g for 5 min, recovered in 4 mL YPD for 4.5–5 h at 30 °C, and distributed over YPD-G418 plates (approximately 100 µL per plate). Transformation plates were photographed and colonies were counted from plate images using OpenCFU software^42^. The first independent construction produced 18,355 HS1-window, 12,858 HS2-window, and 13,704 HS3-window colonies. The second produced 69,458, 29,994, and 31,642 colonies, respectively. Colonies from each regional pool were harvested independently by sequentially washing five plates with 5 mL YPD and pooling the suspensions. Cell suspensions were stored at -70 °C after adding glycerol to 25% of the total volume.

### Dose–response analyses

DMS screens were performed at drug concentrations that inhibit the growth rate (µ) of the WT strain by 50% (IC_50_). To determine these concentrations, dose–response (DR) analyses were first performed in 200 µL volumes (pilot DR) to determine the drug concentration range to test in 10 mL volumes (DMS DR). For the pilot DR, an overnight SC preculture of the WT strain was diluted to OD_600_ 0.15 in 200 µL of SC with a two-fold drug dilution series from 8 µg/mL to 0.0078 µg/mL, and a no-drug control (12 wells), in duplicate, in a flat-bottom polystyrene 96-well plate. The OD_600_ was measured every 15 minutes for 24 h in a BioTek Epoch 2 Microplate Spectrophotometer (Agilent). The maximum derivative growth rate (µ) in each condition was calculated from the OD_600_ time series as follows. For each well, the local slope between consecutive OD_600_ measurements of the growth curve was calculated as:

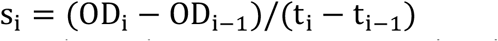

where s_i_ is the local derivative growth rate between two consecutive time points, OD_i_ and OD_i −1_ are consecutive OD_600_ measurements, and t_i_ −t_i-−1_ is the time interval in hours. Because pilot DR measurements were taken every 15 min, this is equivalent to multiplying the OD_600_ difference between adjacent measurements by 4 to express the rate per hour:

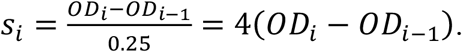

For each well, µ was then defined as the median of the five largest local slopes:

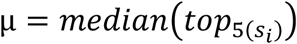

This approach estimates the maximal growth rate during the steepest part of the growth curve while reducing the influence of single noisy measurements. Next, for each drug concentration, growth-rate inhibition was calculated relative to the matched no-drug control:

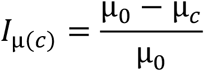

where *I*_μ(*c*)_ is the inhibition coefficient at drug concentration c, μ_*c*_ is the derivative growth rate at concentration c, and μ_0_ is the derivative growth rate of the no-drug control. The resulting coefficients were plotted as a function of drug concentration. The relationship between drug concentration and growth-rate inhibition was fitted with a Hill function:

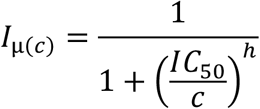

where IC_50_ is the concentration at which the fitted growth-rate inhibition equals 0.5, and h is the Hill coefficient describing the steepness of the dose-response curve (**Supplementary Figure S1A**).

Pilot DR curves were then used to determine a drug concentration range to test in the DMS DR analysis. Cells were treated the same as in the pilot DR analysis, except that they were grown in 10 mL SC volumes and OD_600_ measurements were taken after 3, 6, 8, 10, 12 and 24 h. µ and IC values were calculated in the same way (**Supplementary Figure S1B**). Based on these DMS DR curves, we used the following IC_50_ concentrations in our DMS screens: 0.0176 µg/mL CAS, 0.0286 µg/mL MCF, 0.0711 µg/mL AND, 0.0444 µg/mL RZF, and 0.0985 µg/mL IBX.

### Pooled competition and sampling of DMS screens

The six library pools were labeled HS1_LIB1, HS2_LIB1, HS3_LIB1, HS1_LIB2, HS2_LIB2, and HS3_LIB2 and were screened separately. A frozen aliquot of each pool was inoculated into a 5-mL YPD volume and grown for 12 h at 30 °C with shaking. Precultures were adjusted to OD_600_ 0.30 in 2×SC and mixed 1:1 with 2×drug-containing MilliQ water to obtain 10 mL at OD_600_ 0.15 in 1× SC. Three input samples (t_0_) were collected as pellets containing approximately the same biomass as an output (t_1_, t_2_, t_3_ and t_4_) sample (8 mL at OD_600_ 1.2; 9.6 OD units). Cultures were incubated at 30 °C with shaking and monitored individually. When a culture reached OD600 ≥1.2, a portion was used to inoculate a fresh culture containing the same drug concentration and cell concentration of OD_600_ 0.15. 8 mL of the remaining culture were pelleted and frozen. This OD-controlled transfer was repeated for four passages total, each representing an approximately eightfold expansion (∼3 generations), for a total of approximately 12 generations (t_4_). The t_3_ pellets (9 generations) were used as the post-selection samples.

### Genomic DNA extraction, amplicon library preparation and sequencing

Genomic DNA was extracted from frozen t0 and t3 pellets (1.2 OD_600_) with the MasterPure Yeast DNA Purification Kit (Biosearch Technologies/LGC) according to the manufacturer’s protocol. DNA concentration and purity were measured using a Thermo Scientific NanoDrop 2000c spectrophotometer. To reduce the genomic-DNA bottleneck, each sample was amplified in three parallel 50-µL PCRs containing 500 ng genomic DNA per reaction (1.5 µg total per sample), 10 µL 5× KAPA Fidelity buffer, 1.5 µL 10 mM dNTPs, 1.5 µL each 10 µM primer, 1 µL KAPA polymerase, and water to 50 µL. Cycling was 95 °C for 5 min; 7 cycles of 98 °C for 20 s, 54 °C for 15 s, and 72 °C for 20 s; 13 cycles of 98 °C for 20 s, 72 °C for 15 s, and 72 °C for 20 s; and 72 °C for 3 min. Separate primer pairs distinguished the two independently constructed pools for each region: CLOP372-A8/H8 and CLOP372-B8/G8 for HS1; CLOP372-C8/B9 and CLOP372-D8/A9 for HS2/TM8; and CLOP372-E8/D9 and CLOP372-F8/C9 for HS3 (**Supplementary Table S4**). Parallel PCR products were pooled, purified at a 1:1 sample-to-SPRI-bead ratio, and eluted in 40 µL.

Purified PCR1 products were diluted for the indexing PCR. Each 20-µL index PCR contained 2 µL diluted PCR1 product, 4 µL 5× KAPA Fidelity buffer, 0.6 µL 10 mM dNTPs, 3 µL each 2 µM CLOP224 and CLOP225 row-column index primer (**Supplementary Table S4**), 0.13 µL KAPA polymerase, and 7.27 µL nuclease-free water. Cycling was 98 °C for 3 min; 18 cycles of 98 °C for 30 s, 61 °C for 15 s, and 72 °C for 35 s; and 72 °C for 1 min. Indexed amplicons were SPRI-purified and eluted in 20 µL. Indexed amplicons were prepared for high-throughput sequencing using an i5 and i7 dual-index scheme (**Supplementary Table S4**). The libraries were sequenced at the Plateforme d’Analyses Génomiques (Institut de Biologie Intégrative et des Systèmes (IBIS), Université Laval, Quebec, Canada) on an AVITI instrument (Element Biosciences) using a paired-end 300-base kit following the manufacturer’s instructions. Reads were demultiplexed by sample indexes and assigned to the expected regional library design.

### Deep sequencing data analysis

#### Read preprocessing, gyōza configuration and selection coefficient calculation

Demultiplexed paired-end FASTQ files were processed separately for each library (two per HS- region, 6 total) and split by synonymous barcode (4 per library) for replication quality analysis (see below). Briefly, the hotspot-specific barcode-adjacent 20 nt anchor sequence was identified, the internal synonymous barcode codon extracted, and paired R1/R2 reads were assigned to barcode-specific FASTQ files. Reads requiring more than three anchor edits, including mismatches and/or indels, lacking a detectable anchor, or carrying an unexpected barcode codon were excluded from downstream analyses. Barcode-resolved FASTQ files were analyzed with gyōza v1.2.1^15^, using the codon-based workflow described by Durand *et al*.^15^ but adapted for the barcode-resolved and independently constructed library design used here. The constant sequences supplied to gyōza upstream and downstream of the diversified region were set to 15 nt, which reduced loss of reads due to primer-proximal sequencing errors while retaining region-specific trimming. Gyōza was configured with *Fragment* (HS1, HS2 or HS3), *Barcode* (internal synonymous codon linked to each DMS fragment: for HS1: TCA, TCC, TCG, or TCT encoded serine at position 603; for HS3: GGA, GGC, GGG, or GGT encoded glycine at position 712; and for HS2: CGA, CGC, CGG, or CGT encoded arginine at position 1381), and *Library* (LIB1 and LIB2 for the two independently constructed libraries respectively) as sample attributes and *Condition* (CAS, MCF, RZF, AND, IBX, no_drug for t_3_ samples or CTRL for t_0_ input samples) as the screening attribute. The Library field specified the t_0_–t_3_ pairing for each screen. Only codon– barcode estimates with ≥50 reads in every input (t_0_) replicate were retained for downstream aggregation. Selection coefficients were normalized by the number of generations (normalize_with_gen = True), with t_3_ samples assigned nine generations. For the final heatmaps, estimates for each amino-acid substitution were averaged within each independently constructed library, and the library means were then averaged with equal weight.

Variant read counts were converted to allele frequencies as:

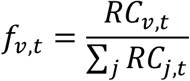

where *f*_*v,t*_ is the allele frequency of variant *v* at time point *t, RC*_*v,t*_ is the read count of variant *v* at time point *t*, and ∑_*j*_ *RC*_*j,t*_ is the total read count across all variants *j* in that sample. For each variant, the change in allele frequency between t_0_ and t_3_ was calculated as:

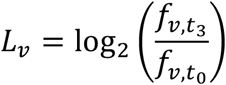

where *L*_*v*_ is the log2 fold-change in allele frequency of variant *v* between t_0_ and t_3_. Selection coefficients were normalized to the number of generations and corrected using the median log_2_ fold-change of silent variants, as implemented in gyōza:

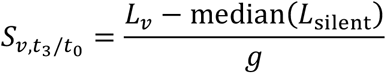

where 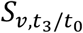 is the generation-normalized selection coefficient of variant *v, L*_silent_ represents the log_2_ fold-change values of synonymous variants used for normalization, and *g* = *t* is the number of generations between t_0_ and t_3_. Positive values indicate enrichment between t_0_ and t_3_, whereas negative values indicate depletion; both are defined relative to synonymous variants. Under drug selection, positive scores indicate a relative growth advantage consistent with resistance. In the no-drug condition, negative scores indicate reduced fitness in the absence of drug.

### DMS replication quality analyses

We evaluated the reproducibility of the DMS screens in four ways.

First, barcode representation was assessed from the demultiplexed FASTQ files before downstream interpretation of selection coefficients. For each targeted region and each independently constructed library, read counts were summed across the barcode-resolved FASTQ files corresponding to each synonymous barcode. The fraction assigned to each expected barcode was calculated as:

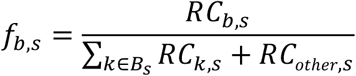

where *f*_*b,s*_ is the fraction of reads assigned to barcode *b* in sample group *s, RC*_*b,s*_ is the corresponding read count, *B*_*s*_ is the set of expected synonymous barcodes for that fragment, *k* indexes the expected barcode classes, and *RC*_other,*s*_ is the number of reads that passed the fragment-specific parsing step but were not assigned to one of the expected barcode bins. These fractions were plotted separately for HS1, HS3, and HS2, with separate rows for LIB1 and LIB2 (**Supplementary Figure S2**).

Second, library-level reproducibility was assessed by comparing selection coefficients for matched variants between independently constructed libraries. For each HS-region, condition, and synonymous barcode, variant scores from LIB1 and LIB2 were merged by the downstream nucleotide variant identifier. Pearson correlation coefficients were calculated from the paired selection coefficients:

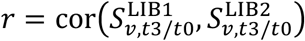

and the coefficient of determination was calculated as: *R*^2^ = *r*^2^ where *r* is the Pearson correlation coefficient, 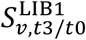 and 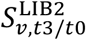 are the generation-normalized selection coefficients for variant *v* estimated from LIB1 and LIB2, respectively, and *t*3/*t*0 denotes the comparison of the selected population at t3 with the matched input population at t0. Points were colored by the synonymous barcode used to resolve the replicate estimate (**Supplementary Figure S3**).

Third, score-difference distributions were used as a complementary measure of reproducibility. For each paired comparison, the difference in selection coefficient was calculated as:

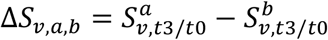

where Δ*S*_*v,a,b*_ is the difference in selection coefficient for variant *v*, and *a* and *b* are two barcode-resolved or library-resolved estimates of the same variant. Two comparison classes were analyzed. “Between barcodes” refers to pairwise comparisons among synonymous barcode groups within the same independently constructed library and condition. “Between libraries” refers to comparisons between LIB1 and LIB2 for the same condition and barcode. Boxplots show the distribution of Δ*S*_*v,a,b*_, with individual variant-level differences overlaid (**Supplementary Figure S4**).

Fourth, the effect of read depth on reproducibility was evaluated by recalculating Pearson correlations after applying increasingly stringent read count thresholds. For each paired estimate, the read count used for thresholding was the smaller read count of the two observations, so that a variant was retained only when both estimates were supported at or above the threshold:

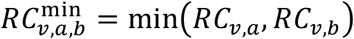

At each threshold *τ*, the retained variant set was defined as:

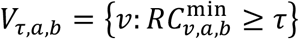

Pearson correlation was then calculated as:

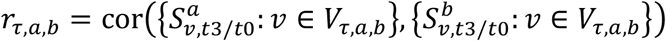

where 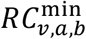 is the minimum read count supporting variant *v* across paired estimates *a* and *b, τ* is the read count threshold, *V*_*τ,a,b*_ is the set of variants retained at read count threshold *τ*, and *r*_*τ,a,b*_is the Pearson correlation coefficient between estimates *a* and *b* at that threshold. Thresholds from 0 to 5000 reads were tested, with denser sampling around the gyōza confidence threshold and the intended sequencing depth. The dashed vertical line indicates the gyōza read count confidence threshold of 50 reads. The dotted vertical line indicates the experimental target coverage of 500 reads per variant per library per barcode. Individual points represent pairwise correlation estimates, lines show the mean correlation across pairwise comparisons within each class, and shaded areas show the standard error of the mean (**Supplementary Figure S5**).

### Mutational Landscape Analyses

To summarize resistance activity across the three targeted *FKS1* regions, selection coefficients were converted into position-level resistance profiles. This analysis was used only as a regional overview; mutation-level effects were analyzed separately from the unscaled nested mean selection coefficients. Repeat-screen conditions were excluded from this overview, and the analysis included no-drug, CAS, MCF, AND, RZF, and IBX.

For each mutation, barcode-level gyōza scores were first summarized within each independently constructed library and then averaged across libraries to obtain the nested mean selection coefficient *S*_*m,c*_, where *m* denotes the amino acid substitution and *c* denotes the condition. Because absolute score ranges differed between drugs and targeted regions, especially for MCF, scores were centered relative to the wild-type amino acid at the same position before comparing position-level activity across conditions:

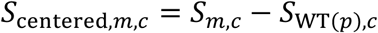

where *S*_centered,*m,c*_ is the WT-centered score for mutation *m* in condition *c, S*_*m,c*_ is the nested mean selection coefficient, and *S*_WT(*p*),*c*_ is the corresponding score of the wild-type amino acid at position *p*. WT centering was used to remove condition-specific and position-specific baseline effects, so that resistance activity reflected the relative effect of an amino acid substitution compared with the wild-type residue at the same position.

Centered scores were then scaled separately for each targeted region and condition using the 95th percentile of the absolute WT-centered scores across all scored substitutions in that region-condition combination:

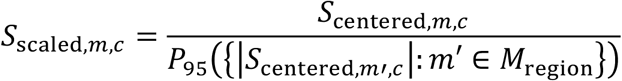

where *S*_scaled,*m,c*_ is the scaled score for mutation *m* in condition *c, M*_region_ is the set of scored substitutions within the corresponding targeted region, and 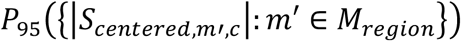 is the 95th percentile of the absolute WT-centered score distribution for all scored substitutions m’ in that targeted region and condition. Scaled scores were clipped between -1 and 1. This scaling converts scores into a relative activity measure within each drug-region combination and prevents weaker global selection in one condition from dominating the regional comparison. Therefore, *S*_scaled_ > 0.5 was interpreted as strong positive activity relative to the robust dynamic range of that condition and region, rather than as a threshold separating signal from noise.

For each amino acid position *p* and condition *c*, we calculated the fraction of substitutions with strong positive scaled activity:

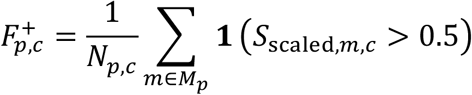

Where 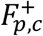 is the fraction of substitutions at position *p* with *S*_scaled_ > 0.5,*M*_*p*_ is the set of scored amino acid substitutions at position *p, N*_*p,c*_ is the number of scored substitutions at that position in condition *c*, and **1**(⋅) equals 1 when the criterion is true and 0 otherwise. For the no-drug condition, depletion was summarized analogously as the fraction of substitutions with strong negative scaled activity:

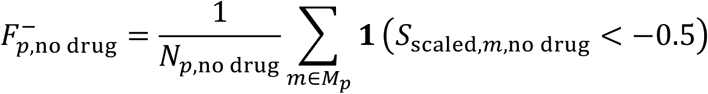

To reduce overinterpretation of isolated single-position fluctuations, position-level fractions were smoothed within each targeted region using a five-position Gaussian kernel with *σ* = 1:

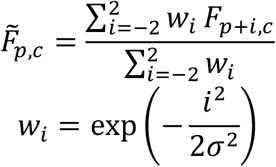

where 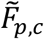 is the smoothed position-level fraction at position *p* and condition *c, F*_*p*+*i,c*_ is the unsmoothed fraction at neighboring position *p* + *i*, and *w*_*i*_ is the Gaussian weight assigned to that neighboring position. Smoothing was performed within each targeted region only and was not allowed to connect non-contiguous DMS fragments. This approach assumes that resistance hotspots represent local clusters of functionally related residues rather than isolated single substitutions.

We defined an echinocandin-resistance region as a near-continuous region in which resistance-associated substitutions recurred across multiple amino acid changes and across multiple echinocandins. Because MCF showed a lower dynamic range and weaker directional signal than the other echinocandin conditions, ECN-resistance region calling was based on CAS, AND, and RZF, while MCF was retained in the plot for comparison. A position was called ECN-active when the mean smoothed fraction across CAS, AND, and RZF was at least 0.15, and at least two of these three echinocandins had 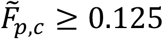. IBX-resistance regions were defined independently as positions where the smoothed IBX fraction was at least 0.15. Resistance regions were then defined, allowing internal gaps of up to two residues. This gap rule was used to avoid fragmenting a broader local activity region when one or two intervening positions fell below the threshold, while preventing more distant activity peaks from being merged into a single hotspot. Historical echinocandin hotspot intervals were plotted as reference regions but were not used to calculate the DMS-defined resistance regions.

Mutation-level fitness heatmaps were generated from the generation-normalized gyōza selection coefficients after confidence filtering. Barcode-level estimates were first averaged within each independently constructed library for each mutation and condition:

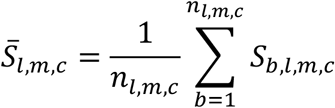

where 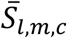 is the library-level mean selection coefficient for library *l*, mutation *m*, and condition *c, S*_*b,l,m,c*_ is the barcode-level selection coefficient, and *n*_*l,m,c*_ is the number of barcode-resolved estimates available for that mutation, library, and condition. Library-level means were then averaged across independently constructed libraries:

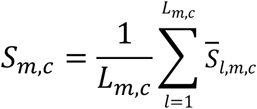

where *S*_*m,c*_ is the nested mean selection coefficient used for heatmap visualization and *L*_*m,c*_ is the number of independently constructed libraries with a score for mutation *m* in condition *c*. This nested averaging gives equal weight to independently constructed libraries and prevents synonymous barcode estimates from being treated as independent biological replicates.

For the main mutation-level heatmaps in Figure 2, the plotted regions were restricted to the union of the ECN- and IBX-resistance regions defined from the position-level activity analysis, extended by seven amino acid positions upstream and downstream and clipped to the experimentally targeted DMS region. This yielded plotting windows of positions 624-656 for HS1, 679-707 for HS3, and 1346-1380 for HS2. Full heatmaps across the complete targeted regions are shown in **Supplementary Figure S6**. Wild-type residues were indicated by black dots on light grey tiles, unscored substitutions were shown as dark grey tiles, and priority substitutions from the FungAMR and Gifford analyses were outlined in green.

Differential selection heatmaps were generated by subtracting nested mean scores between conditions:

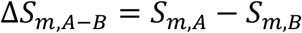

where Δ*S*_*m,A*−*B*_ is the differential selection coefficient for mutation *m, S*_*m,A*_ is the nested mean selection coefficient in condition *A*, and *S*_*m,B*_ is the nested mean selection coefficient in condition *B*. For Figure 2B, condition *A* was AND and condition *B* was IBX. Positive values therefore indicate higher selection coefficients under AND than IBX, whereas negative values indicate higher coefficients under IBX than AND. Equivalent comparisons of CAS, MCF, RZF, and AND versus IBX were generated for **Supplementary Figure S7**.

For each position *p*, differential effects were summarized in two ways. First, the mean differential selection coefficient was calculated across all scored substitutions at that position:

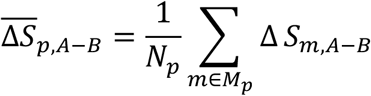

where *M*_*p*_ is the set of scored amino acid substitutions at position *p*, and *N*_*p*_ is the number of scored substitutions at that position. Second, the number of substitutions with strong differential effects was calculated as:

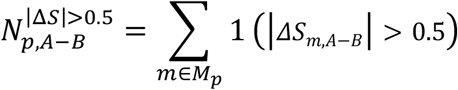

where 1(*l*Δ*S*_*m,A*−*B*_ *l >* 0.*5*)equals 1 when the absolute differential selection coefficient of mutation *m* exceeds 0.5 and equals 0 otherwise. These summaries were plotted below the differential heatmaps, with the mean differential score shown as a line and the number of substitutions with |Δ*S*| *>* 0.5 shown as bars.

### Prioritization of substitutions of interest

To define a focused set of priority substitutions, we combined two complementary sources of evidence: the *C. auris* whole-genome sequencing dataset of Gifford *et al*.^6^, which represents the largest genome dataset assembled for a fungal pathogen to date, and the curated FungAMR database of antifungal resistance-associated variants^16^.

Fks1 variation frequencies from the Gifford *et al*.^6^ *C. auris* genome dataset were mapped on an alignment of *C. auris* Fks1 (type strain B8441) and *S. cerevisiae* Fks1 sequences. Only substitutions falling within the *C. auris* regions corresponding to the *S. cerevisiae* DMS-targeted regions were retained, and since our DMS libraries only quantified amino acid substitutions, deletions and insertions were excluded. For each retained *C. auris* substitution, overall prevalence across the analyzed clades was calculated and plotted in **Supplementary Figure S8**. Substitutions with an overall prevalence of at least 0.05% (minimum 6 out of 12 644 analyzed genomes) were considered recurrent *C. auris* population variants.

In parallel, FungAMR^16^ entries were filtered for Fks1 substitutions associated with echinocandin-class drugs. Each mutation was ranked by the number of independent source identifiers reporting that substitution. Ties were resolved by confidence score, number of species, and total number of supporting entries. The ten most frequently reported exact substitutions were retained as literature-prioritized mutations, shown in **Supplementary Table S2**.

The final set of substitutions of interest was defined as the union of the ten most frequently reported exact FungAMR^16^ substitutions and recurrent *C. auris* substitutions detected at ≥0.05% prevalence in the Gifford *et al*. dataset^6^. These substitutions were marked on both the *S. cerevisiae* DMS fitness heatmaps in **Figure 2** and **Supplementary Figures S6-7** and the *C. auris* prevalence heatmap in **Supplementary Figure S8**.

### Construction of individual *C. auris* mutants

To construct allelic variants, plasmid pJMR19 from Doorley *et al*.^43^ was digested with SapI and ligated using the respective duplexed oligos as mentioned in **Supplementary Table S4**. The top and bottom guide oligos were duplexed in a 50 μL final volume and 2 μM final concentration. For ligation, 40 ng of the pJMR19 SapI digest were mixed with 100 nM final concentration of the duplexed oligos in a final reaction volume of 10 μL, using the T4 DNA ligase enzyme from NEB, according to the manufacturer’s instructions. The plasmids used in this study were propagated in *E. coli* TOP10F’ chemically competent cells. Bacterial transformations were carried out by heat shock at 42 °C for 45 s using 30 μL of competent cells, and subsequent cooling on ice for 2 minutes. The transformants were selected on solid LB (Sigma, Fisher Scientific) agar supplemented with ampicillin (100 μg/mL). Plasmid DNA was isolated using the NucleoSpin Plasmid kit (MN) according to the manufacturer’s instructions, and PCR products were purified using the NucleoSpin Gel and PCR Clean-up XS kit (MN). For enzymatic digestions, the final volume of the reaction was 50 μL with 1 μL of each restriction enzyme and the amount of DNA ranged from 2 to 10 μg depending on the desired amount. Molecular analyses for plasmid construction were performed in SnapGene 7.0.

Strains were constructed following a previously published approach^44,45^. In short, a single colony of *C. auris* strain B8441 was inoculated in liquid YPD and grown overnight at 37 °C in a shaking incubator. The preculture was diluted in 50 mL YPD in a conical flask to an OD_600_ of 0.4 and grown until the OD_600_ reached a range of 1.6 to 2.2 (approximately 3-4 hours). The cells were collected (5 minutes at 3 273 x g), resuspended in 10 mL of transformation buffer [10 mM Tris-HCl, 1 mM EDTA (VWR) and 100 mM LiOAc (Sigma)] and shaken at 37 °C, 150 rpm for 1 hour. 250 μL of 1 M DTT (VWR) was added, and the cells were incubated for an additional 30 minutes. Cells were washed twice (5 minutes at 5 000 x g at 4 °C), first with 25 mL ice-cold dH_2_O and then with 5 mL ice-cold 1 M sorbitol (Sigma). The supernatant was removed carefully, and the pellet was resuspended in 200 μL of ice-cold 1 M sorbitol. 40 μL of the competent cells was mixed with the transformation mixture and transferred to a 2 mm electroporation cuvette (Pulsestar, Westburg). A single pulse was given at 1.8 kV, 200 Ω, 25 μF, and the transformation mixture was immediately transferred to 2 mL YPD in test tubes following incubation for 4 hours at 37 °C, 150 rpm. The cells were collected by centrifugation for 5 minutes at 5 000 x g, resuspended in 100 μL YPD and plated on YPD agar containing 200 mg/mL of nourseothricin (Jena bioscience). Transformants appeared after two to three days of incubation at 37 °C. For point mutant constructs, the transformation mixture comprised 5 μg of the corresponding pJMR19 construct and 5 μg of the dDNA, which was amplified from a gblock or mutant strain using primers displayed in Supplementary Table S4. The locus of interest of each transformant was PCR amplified and sequenced to verify correct transformants using primers displayed in **Supplementary Table S4**. Correct transformants were recycled by growing sequentially on non-selective YPD medium until loss of nourseothricin resistance was apparent by replicate plating on YPD and YPD+NTC.

### Susceptibility testing of individual mutants

Antifungal susceptibility testing was done by the EUCAST reference method^46^ with minor adaptations. Briefly, a twofold dilution range of drug from 2 µg/mL to 0.008 µg/mL was prepared in a total volume of 200 µL RPMI-MOPS (pH 7, 2% glucose, 1% DMSO) medium with approximately 200 cells (based on OD_600nm_ and serial dilution) in a round-bottom 96-well polystyrene microtiter plate (Greiner). Plates were incubated at 37 °C for 48 hours, and growth was assessed spectrophotometrically (OD_600_) using a Synergy™ H1 microplate reader (BioTek). The growth cut-off of all MIC values from the broth dilution assay (BDA) was 50% growth compared to the drug-free control.

### Fks1-inhibitor structural modeling and binding-pose selection

#### Model preparation

Molecular models were prepared from the *S. cerevisiae* Fks1 cryo-EM structure in complex with CAS, nascent glucan, membrane-associated lipids, Rho1, and Gsr1; PDB 9PE1^7^. This structure was selected because it provides experimentally determined positions for CAS and the elongated glucan chain within the translocation pathway. The initial complex was prepared using the CHARMM-GUI PDB Reader and Membrane Builder workflow, retaining Fks1 (chain 1), Rho1 (chain 2), Gsr1 (chain 3), glucan, and the resolved 1,2-distearoyl-sn-glycero-3-phosphoethanolamine (DSPE)^47-49^. Glycosylations at Fks1 residues 1845 and 1849 and disulfide bonds between residues 658-669 and 1328-1345 were retained. Protonation states were assigned at pH 7 using CHARMM-GUI, including the nonstandard patches ASPP135, ASPP180, ASPP684, ASPP1084, ASPP1416, LYSN632, and ASPP691 in segment PROA, and CYM21 in PROB. Default terminal patches were otherwise applied. Missing Fks1 residues 242-265, 594-596, 799-803, 902-913, and 1715-1720 were modeled using MODELLER through CHARMM-GUI. The complex was oriented relative to the membrane using PPM 2.0, as implemented in CHARMM-GUI, with all three protein chains^50^.

#### Sterol replacement

Cholesteryl hemisuccinate (CHS) molecules resolved in PDB 9PE1 were replaced with ergosterol using a custom Python script and RDKit^51^. The resulting coordinates were processed with the CHARMM-GUI PDB Reader and Manipulator and combined with the prepared Fks1 system using CHARMM version 50^52^. This procedure preserved the experimentally observed positions of the membrane-associated sterols while replacing CHS with ergosterol, the major sterol of fungal plasma membranes.

#### Ligand building and parameterization

The prepared Fks1-glucan-ergosterol model served as the common structural template for the five inhibitors. For CAS, the experimentally resolved ligand coordinates from PDB 9PE1 were retained, and the corresponding PDB segments were combined into a single ligand segment. MCF, AND, and RZF were constructed using the PyMOL Builder tool and initially positioned by structural superposition onto the CAS structure. IBX was modeled using the enfumafungin-bound Fks1 cryo-EM structure PDB 9WZU^11^. The Fks1 structures from PDB 9WZU and 9PE1 were aligned using the regions encompassing the three canonical resistance hotspots, and IBX was built and positioned using the enfumafungin binding geometry as a template. Although no glucan was resolved in the enfumafungin-bound cryo-EM structure, the nascent glucan chain from the CAS-bound structure (PDB 9PE1) was retained in the IBX model. This approach allowed us to investigate IBX binding in the context of a glucan-containing Fks1 complex, representing a catalytically engaged state of the enzyme, and to assess potential interactions between IBX and nascent glucan. It also provided a consistent structural framework for comparing the binding modes of echinocandins and IBX.

Each ligand was processed through the CHARMM-GUI Ligand Reader and Modeler to generate molecular topology, coordinate, and force-field parameter files^53^. Ligand parameters were assigned using the CHARMM General Force Field (CGenFF) program version 4.0, with CGenFF topology and parameter files version 5.0^54^. The parameterized ligands were merged individually with the prepared Fks1 system using CHARMM, generating five Fks1-glucan-inhibitor models.

#### Simulated annealing

Each model was refined by simulated annealing in CHARMM using the generalized Born with simple switching (GBSW) implicit-solvent model with an implicit membrane representation^55,56^. The simulations used CHARMM-GUI-generated PSF and coordinate files together with the corresponding protein, lipid, carbohydrate, and ligand topology and parameter files. Atomic radii were assigned using radii_prot_na.str and radius_gbsw.str, with default radii applied to other atoms. The GBSW implicit-membrane parameters included a membrane thickness of 30 Å, a membrane switching length of 2.5 Å, a surface-tension coefficient of 0.04 kcal mol^−1^ Å^− 2^, a salt concentration of 0.15 M, a maximum Born-radius integration distance of 20 Å, and 38 angular integration points.

For each inhibitor, 20 independent simulation replicas were performed using different initial random seeds, each comprising 20 consecutive simulated-annealing cycles. Before the first cycle, the initial structure was minimized using 200 steps of steepest descent (SD) followed by 200 steps of adopted-basis Newton–Raphson (ABNR) minimization, and then equilibrated for 5,000 steps of Langevin dynamics at 303.15 K. Each annealing cycle consisted of two successive 5,000-step Langevin dynamics segments at 500 K and 303.15 K, respectively. All dynamics were performed with a 2 fs timestep, a Langevin friction coefficient of 5 ps^−1^, and SHAKE constraints on bonds involving hydrogen atoms. Each temperature segment lasted 10 ps, yielding 20 ps of dynamics per annealing cycle. Following each cycle, the system was minimized using 250 steps of SD followed by 250 steps of ABNR minimization. Each subsequent cycle continued from the preceding cycle rather than restarting from the initial structure. The minimized coordinates from each cycle were saved for structural and energetic analyses, yielding 400 candidate binding poses per inhibitor.

#### Binding-pose evaluation and ranking

Candidate binding poses were evaluated using a combination of DMS-informed, structural, and energetic criteria. Agreement with the experimental DMS resistance landscapes was assessed from the number of DMS-identified resistance-associated residues in contact with each inhibitor (DMS_residues). Structural agreement with the reference Fks1-glucan geometry from PDB 9PE1 was evaluated using root-mean-square deviation (RMSD) measurements. Candidate poses with a glucan RMSD greater than 5 Å, calculated after alignment of the protein to the reference Fks1– glucan structure, were excluded from the global ranking.

Energetic ranking considered both the ligand internal energy and the ligand binding energy, with more favorable poses defined by lower values. Binding energy was calculated using the GBSW membrane implicit-solvent model and the parameters described above, according to:

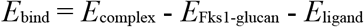

where *E*_complex_ is the energy of the Fks1-glucan-ligand complex, *E*_Fks1-glucan_ is the energy of the Fks1-glucan component, and *E*_ligand_ is the energy of the isolated ligand. These values represent model-dependent binding energy estimates rather than binding free energies.

Candidate poses were ranked by combining the DMS-derived contact criterion, structural deviations, and energetic scores. The top-ranked poses were subsequently examined for their predicted interactions with Fks1 and glucan and for consistency with the available cryo-EM structures.

#### Software

DMS analysis was performed using gyōza v1.2.1^15^. Custom preprocessing and post-gyōza analyses were performed in Python 3.13 using scripts developed and run in Visual Studio Code. Data handling and numerical analyses were performed with pandas and NumPy, and figures were generated with matplotlib and seaborn. Excel workbooks were read and written using openpyxl. Custom scripts were used for barcode-resolved FASTQ splitting, post-gyōza quality-control analyses, generation of mutation-level heatmaps, differential-selection plots, position-level resistance-region plots, and integration of FungAMR^16^ and *C. auris* population-variation data^6^. Figure 1B was created in BioRender, licensed by Van Dijck, P. (2026) https://BioRender.com/epekrdc. MD simulations were performed using CHARMM version 50^52^. Structural analyses and binding-pose ranking were performed using Python (Python v3.12) within a dedicated Conda environment (Conda v25.11.1). Molecular structures were visualized and inspected, and 3D images were produced, with Open-Source PyMOL (PyMOL v3.1.0)^57^. The following Python package versions were used: Biopython v1.86^58^, RDKit v2025.03.6^51^, NumPy v2.4.2^59^, pandas v3.0.3^60^, MDAnalysis v2.10.0^61^, matplotlib v3.10.8^62^.

## Acknowledgments

H.C. was supported by a postdoctoral fellowship from the European Molecular Biology Organization – EMBO (ALTF 1105-2024), and travel grants from le regroupement québécois de recherche sur la fonction, l’ingénierie et les applications des protéines – PROTEO, le Centre de Recherche en Infectiologie – CRI, the Canadian Bioinformatics Hub – CBH, the Genetics Society of America – GSA & the Canadian training program on the Evolution of Fungal Pathogens – EvoFunPath. S.J. & D.S. were supported by PhD fellowships from the Fund for Scientific Research Flanders – FWO (11PRR24N & 11J8122N, respectively). R.D. was supported by an NSERC postdoctoral fellowship. The group of P.V.D. was supported by FWO under the framework of the JPIAMR – Joint Programming Initiative on Antimicrobial Resistance fund (project CycleDrug) and by a C3 grant from the Industrial Research Fund of KU Leuven (C3/22/007). The group of C.R.L. was supported by a Genome Canada and Genome Québec grant (no. 6569). C.R.L. holds the Canada Research Chair in Cellular Systems and Synthetic Biology. The group of P.L. was supported by the Natural Sciences and Engineering Research Council of Canada (NSERC; Grant RGPIN-2022-04721) and by support from Calcul Québec (calculquebec.ca) and the Digital Research Alliance of Canada (alliancecan.ca).

We would like to thank our collaborators Shivaprakash Rudramurthy, Suhail Ahmad, Katrien Lagrou, Alexander Lorenz & Tatsuro Hirayama for sharing *C. auris* isolates, Jeffrey M. Rybak for sharing plasmid pJMR19, and Hugh Gifford and Rhys Farrer for sharing data from Gifford *et al*.^6^.

## Author contributions

Conceptualization: H.C. Methodology: H.C., A.K.D., R.D., P.L. & C.R.L. Software: H.C., R.D. & P.L. Investigation: H.C., N.W., S.J., A.K.D., D.S., C.L.R., B.S. & P.L. Formal analysis: H.C., N.W. & P.L. Resources: H.C., E.M., P.V.D., P.L. & C.R.L. Data curation: H.C., N.W. & B.S. Writing – original draft: H.C. Writing – review and editing: H.C., N.W., S.J., A.K.D., D.S., R.D., B.S., E.M., P.V.D., P.L. & C.R.L. Visualization: H.C., N.W. & P.L. Supervision: H.C., E.M., P.V.D., P.L. & C.R.L. Project administration: H.C., P.L. & C.R.L. Funding acquisition: H.C., P.V.D., P.L. & C.R.L.

## Data and materials availability

Accession numbers for raw sequencing data, repository links for code and strain availability information will be provided upon formal publication.

