## supplementary for "Distinct binding modes shape exploitable asymmetries between echinocandin and triterpenoid resistance landscapes in Fks1"

#### Supplementary Results

##### Annex 1. Barcode- and library-resolved reproducibility of DMS measurements

The barcode fraction analysis showed that the four synonymous barcode classes were recovered in all three HS-regions and in both independently constructed libraries (**Supplementary Figure S2**). Across conditions, expected barcode fractions ranged from 16.1-30.9% in HS1, 15.6-38.8% in HS3, and 15.4-34.2% in HS2. Reads that could not be assigned to an expected barcode class because of an unrecognized flanking anchor, an incomplete barcode, or an unexpected barcode codon were classified as "other". These reads accounted for 0.4–4.7% in HS1, 1.5–5.2% in HS3, and 1.4–3.2% in HS2. These results indicate that barcode splitting did not produce major barcode dropouts and that most barcode-resolved replicates were represented in the input data. HS3 showed somewhat more barcode imbalance than HS1 and HS2, most notably in several IBX-containing samples where one barcode represented a larger fraction of reads, but the imbalance was not severe enough to remove a barcode class from downstream analyses.

LIB1/LIB2 score correlations supported reproducible drug-selected DMS measurements for most conditions (**Supplementary Figure S3**). In HS1, drug-selected conditions showed moderate to strong agreement between independently constructed libraries, with Pearson r values of 0.56 for MCF, 0.67 for CAS, 0.77 for AND, 0.79 for RZF, and 0.80 for IBX. HS2 showed similar reproducibility, with drug-condition correlations ranging from 0.51 for MCF to 0.67 for RZF. HS3 showed weaker library-level correlations overall, with r values of 0.28 for MCF, 0.43 for IBX, 0.48 for AND, 0.49 for RZF, and 0.50 for CAS. This indicates that HS3 contains reproducible signal, but with a lower signal-to-noise ratio than HS1 and HS2, particularly for MCF. The no-drug condition showed low LIB1/LIB2 correlations in all fragments (HS1 r = 0.28, HS3 r = 0.16, HS2 r = 0.15), which is okay, as in the absence of drugs, positive selection for resistance alleles is not expected; most variants are near neutral or depleted, so between-library variation and sampling noise contribute proportionally more to the observed score differences.

The score-difference distributions further supported the absence of systematic biases between barcode or library estimates (**Supplementary Figure S4**). Across all fragments, median score differences were close to zero. Comparisons were performed at the nucleotide-variant level, before averaging synonymous codons or barcodes. Within libraries, selection coefficients were compared for the same nucleotide variant across different barcodes; between libraries, comparisons matched both the nucleotide variant and barcode. The reported values are medians of these pairwise differences in selection coefficient (ΔS). For within-library comparisons between barcodes, the median $\Delta S_{t3/t0}$ was 0.006 for HS1, -0.006 for HS3, and 0.029 for HS2. For between-library comparisons, the median $\Delta S_{t3/t0}$ was -0.007 for HS1, -0.010 for HS3, and 0.027 for HS2. The interquartile ranges were broader for between-library comparisons than for within-library barcode comparisons, as LIB1 and LIB2 represent independently constructed mutant pools and independently screened samples. The distributions were nevertheless centered near zero, indicating no strong global shift between barcodes or between libraries. Wider tails in some no-drug and lower-correlation drug conditions indicate that a subset of variants has less reproducible quantitative scores and should be interpreted with greater caution when effects are subtle or supported by only one replicate class.

The primary analyses used gyōza's high-confidence criterion of ≥50 reads per nucleotide variant in every input ($t_0$) replicate. Higher thresholds were examined to assess the dependence of replicate correlations on read depth (**Supplementary Figure S5**). At 50 reads, median drug-condition correlations were 0.86 for HS1 and 0.83 for HS2; at 500 reads, median drug-condition correlations remained high at 0.70 for HS1 and 0.76 for HS2. HS3 showed a different pattern: the median drug-condition correlation at 50 reads was lower, but increased at higher read-count thresholds, reaching a median of 0.67 at 500 reads. This pattern suggests that HS3 reproducibility was more sensitive to read depth and is consistent with greater measurement noise among low-count variants in this region. At very high read-count thresholds, especially above approximately 2500-5000 reads, several curves became unstable or decreased. This should not be interpreted as evidence that high read count variants are unreliable. Instead, very stringent thresholds retain only a small and biased subset of variants, often those strongly enriched or otherwise highly represented after selection. Correlations at these high thresholds therefore reflect the behavior of a limited subset rather than the reproducibility of the full mutant library. The most relevant range for judging screen quality is around the gyōza confidence threshold and the intended coverage target. In that range, the data support good reproducibility for most drug-selected conditions, especially in HS1 and HS2. Lower correlations in no-drug samples and in some HS3/MCF comparisons indicate reduced quantitative precision for those conditions, but do not undermine the major resistance-enrichment patterns when effects are strong and consistent across barcode and library estimates.

The overall conclusion from these analyses is that the barcode-resolved and independently constructed library design produced reproducible DMS measurements. The strongest and most biologically interpretable signals are those that are consistent across synonymous barcodes and across LIB1/LIB2. Conditions or regions with weaker correlations should be interpreted with more emphasis on replicated, high-confidence substitutions and less emphasis on isolated variants with small effect sizes.

### Supplementary Figures

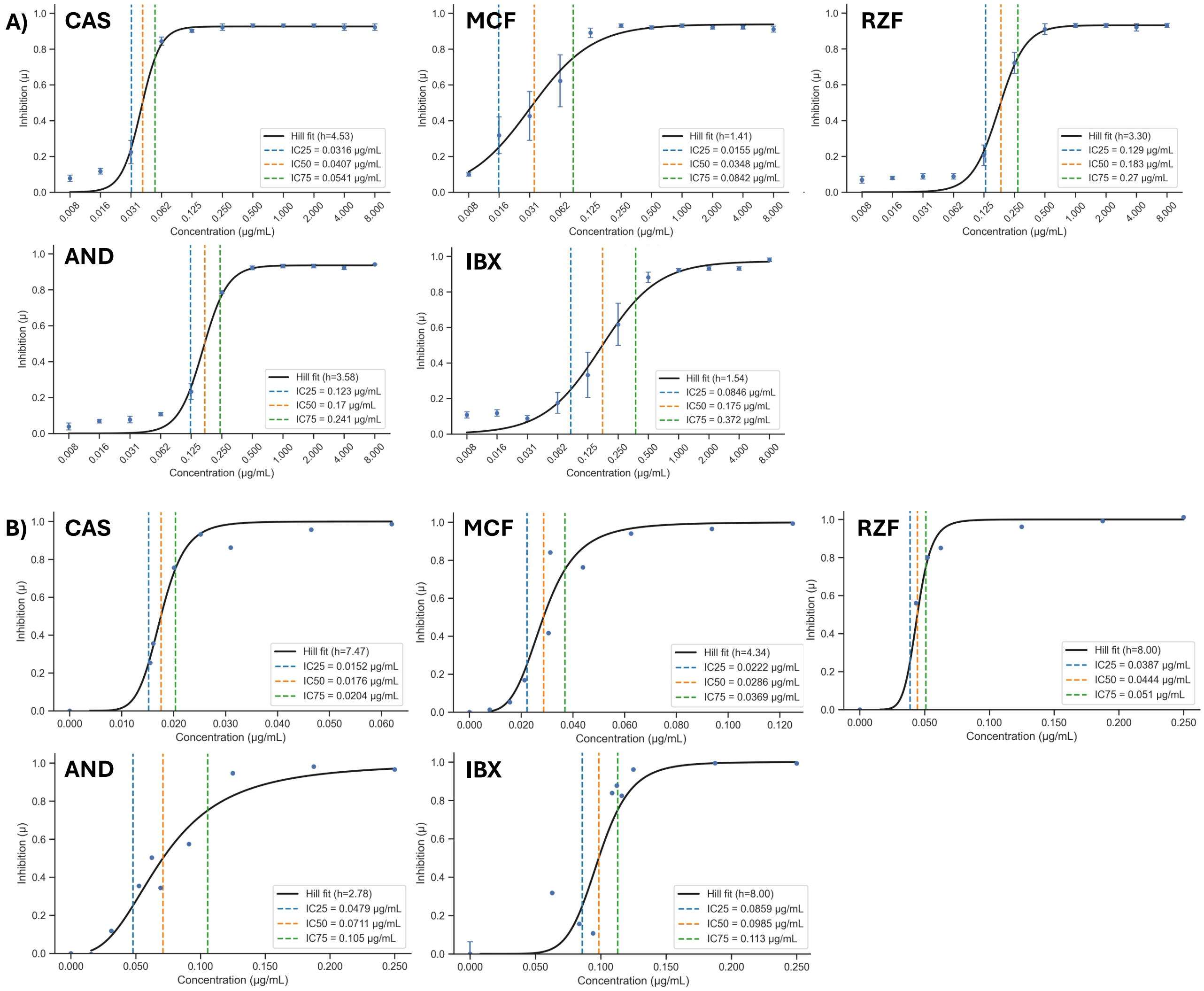


**Figure S1. Dose-response (DR) analyses to determine $IC_{50}$ concentrations for DMS screens.** Pilot DR analysis of the WT strain in 200 μL cultures (**A**) and in 10 mL cultures under the conditions used for DMS screening (**B**). The y-axis shows the growth inhibition coefficient, calculated from the derivative growth rate (μ) of $OD_{600}$ growth curves, relative to the no-drug control. Hill DR curves were fitted to the measured inhibition coefficients as a function of drug concentration. The Hill coefficient describes the steepness of the fitted DR relationship. $IC_x$ is the drug concentration that inhibits μ by x%. $IC_{50}$ values in 10mL were selected as the final drug concentrations for DMS screening.

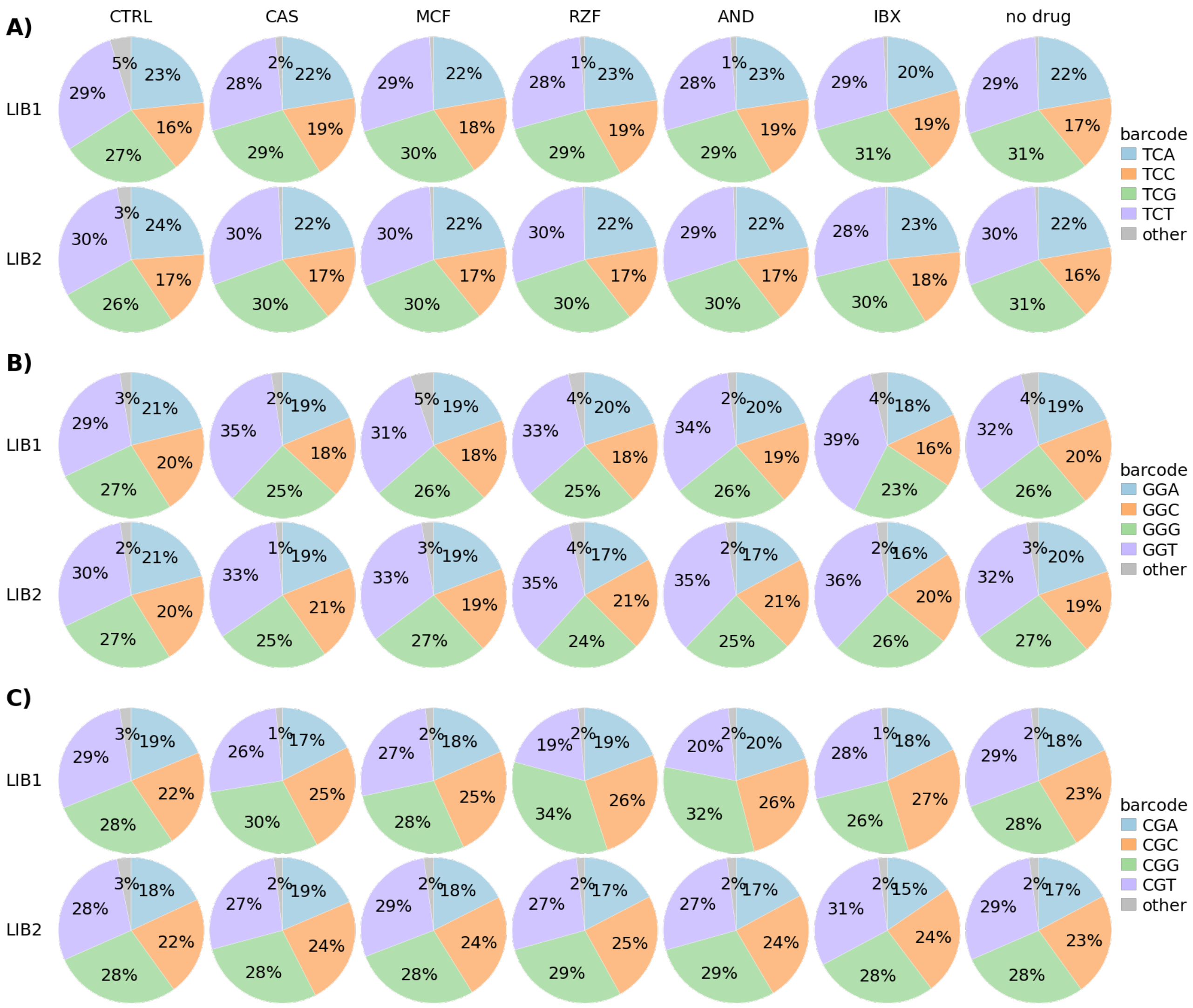


**Figure S2. Barcode representation in demultiplexed DMS libraries.** Barcode fractions are shown for each condition, library, and targeted HS1-region (**A**), HS3-region (**B**) and HS2-region (**C**). Each pie chart shows the fraction of reads assigned to the expected synonymous barcode classes or to the other category. CTRL panels show $t_0$ samples.

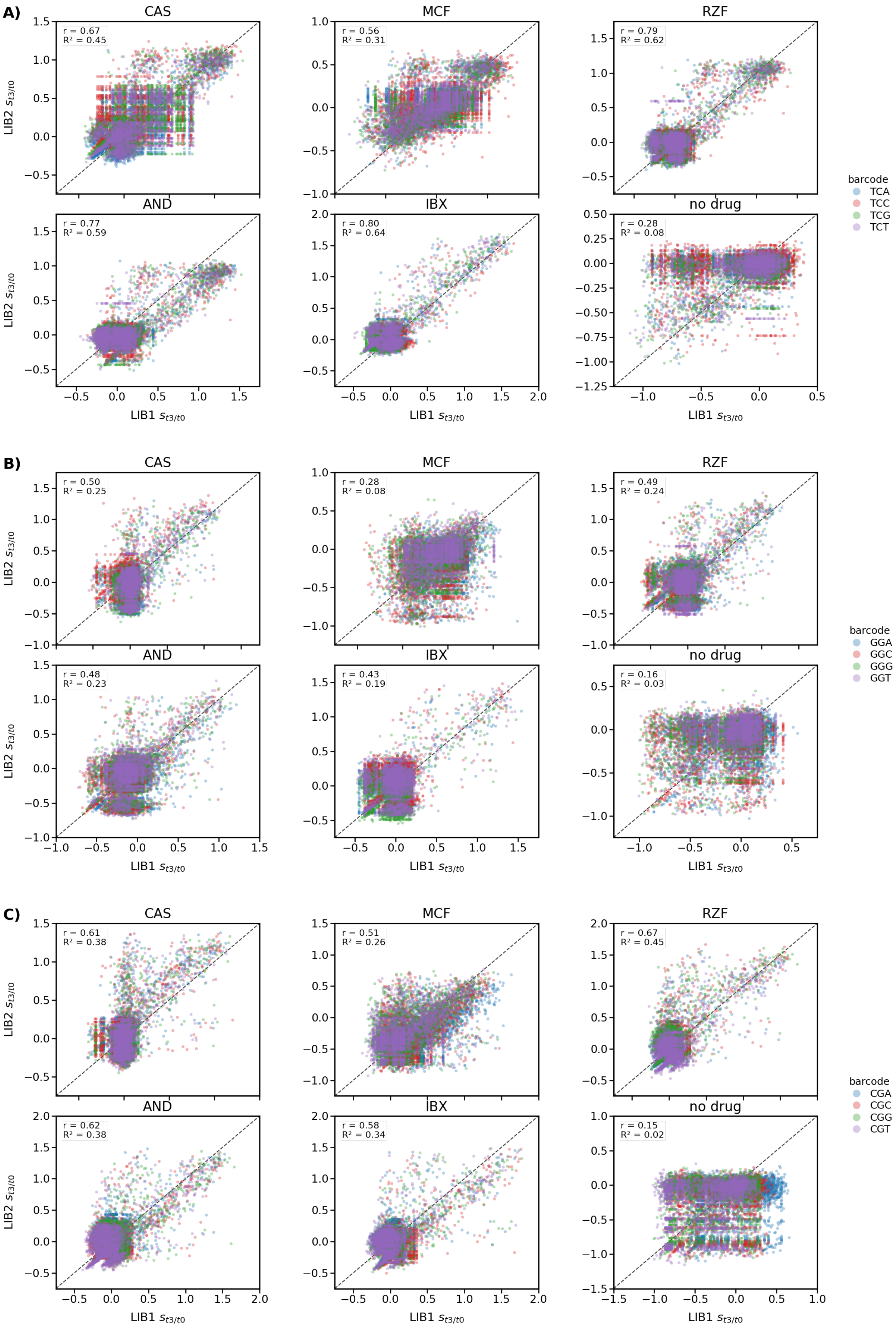

A)
CAS
r = 0.67
R² = 0.45
MCF
r = 0.56
R² = 0.31
RZF
r = 0.79
R² = 0.62
AND
r = 0.77
R² = 0.59
IBX
r = 0.80
R² = 0.64
no drug
r = 0.28
R² = 0.08
barcode
TCA
TCC
TCG
TCT
B)
CAS
r = 0.50
R² = 0.25
MCF
r = 0.28
R² = 0.08
RZF
r = 0.49
R² = 0.24
AND
r = 0.48
R² = 0.23
IBX
r = 0.43
R² = 0.19
no drug
r = 0.16
R² = 0.03
barcode
GGA
GGC
GGG
GGT
C)
CAS
r = 0.61
R² = 0.38
MCF
r = 0.51
R² = 0.26
RZF
r = 0.67
R² = 0.45
AND
r = 0.62
R² = 0.38
IBX
r = 0.58
R² = 0.34
no drug
r = 0.15
R² = 0.02
barcode
CGA
CGC
CGG
CGT
LIB2 $s_{t3/t0}$
LIB1 $s_{t3/t0}$

**Figure S3. Correlation of selection coefficients between independently constructed libraries and among barcodes.** Each point represents one variant $S_{t3/t0}$ for a specific synonymous barcode, coloured by barcode identity, for all conditions in the HS1-region (**A**), HS3-region (**B**) and HS2-region (**C**) libraries. Dashed diagonal lines indicate equal scores in LIB1 and LIB2. Pearson r and $R^2$ values are shown in each panel.

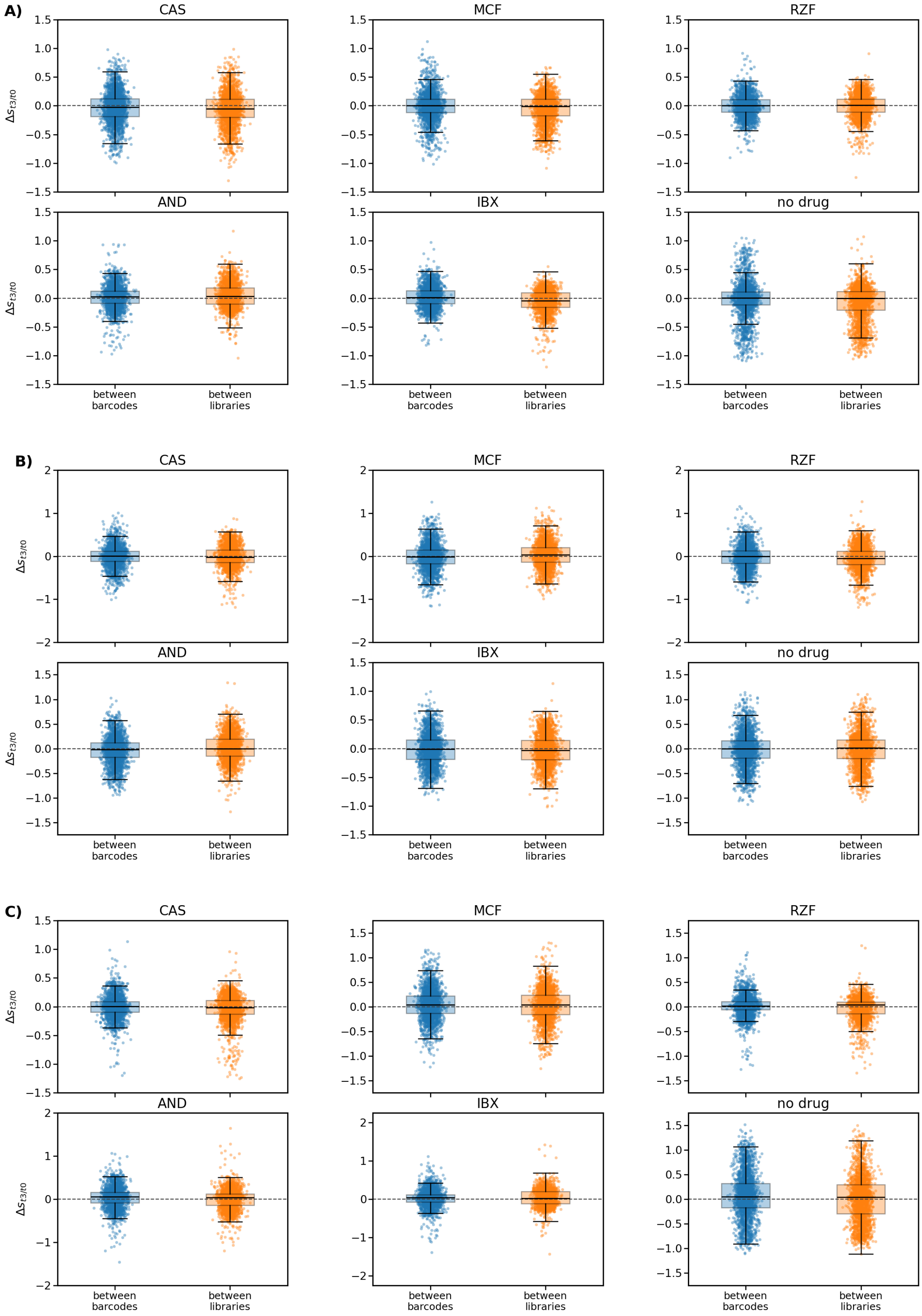
A)
B)
C)
CAS
MCF
RZF
AND
IBX
no drug
$\Delta s_{t3/t0}$
between barcodes
between libraries

**Figure S4. Distribution of selection-score differences between barcode and library estimates.** Distributions of $\Delta S_{t3/t0}$ are shown for barcode-resolved and library-resolved replicate comparisons for HS1-region (**A**), HS3-region (**B**) and HS2-region (**C**). "Between barcodes" compares synonymous barcode estimates within the same independently constructed library, whereas "between libraries" compares LIB1 and LIB2 estimates for the same barcode.

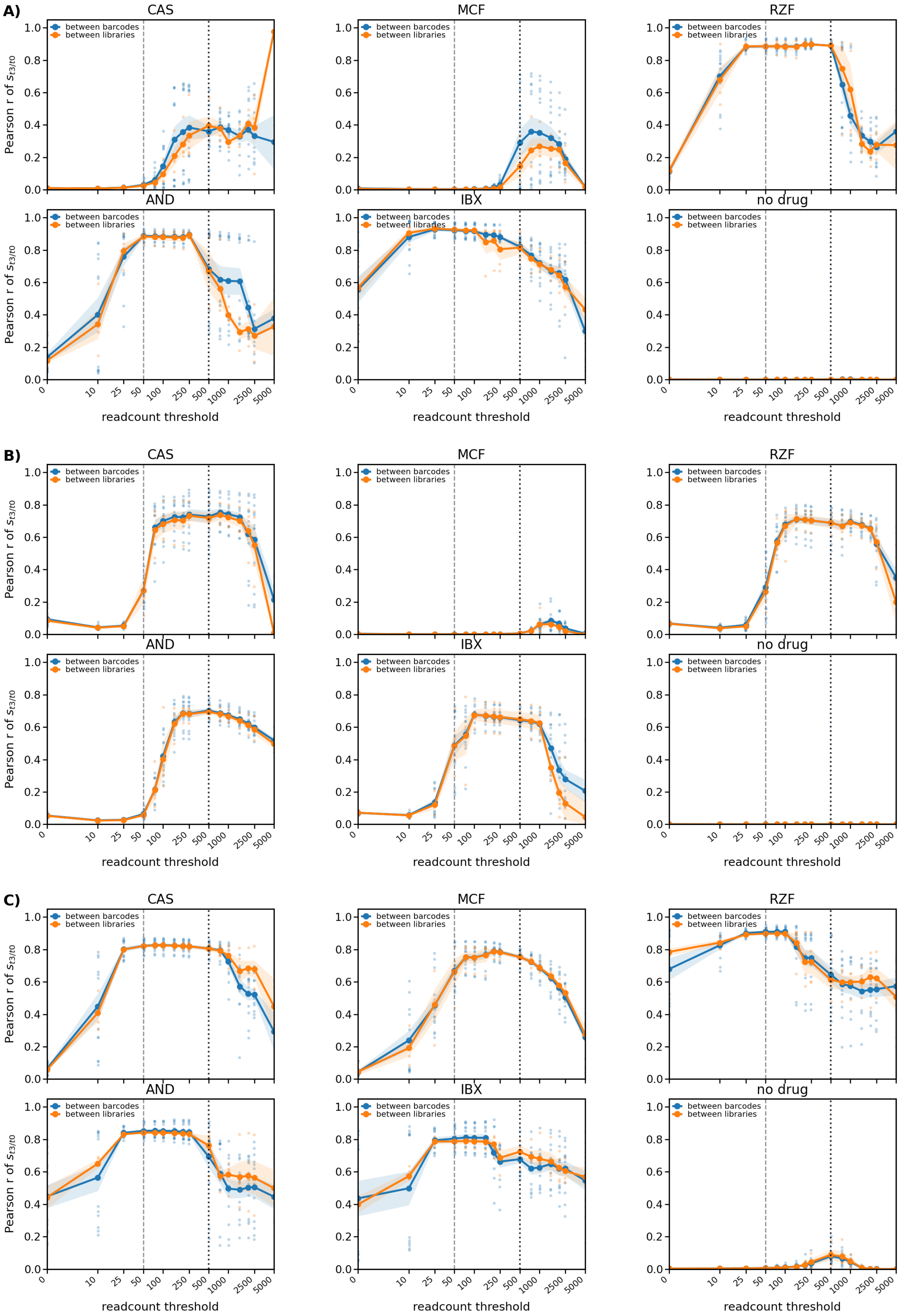
A)
B)
C)
CAS
MCF
RZF
AND
IBX
no drug
between barcodes
between libraries
Pearson r of $s_{t3/t0}$
readcount threshold
0
10
25
50
100
250
500
1000
2500
5000
0.0
0.2
0.4
0.6
0.8
1.0

**Figure S5. Dependence of reproducibility on read count threshold.** Pearson correlations between replicate selection coefficient estimates were recalculated after applying increasing minimum read count thresholds for the HS1-region (**A**), HS3-region (**B**) and HS2-region (**C**). The x-axis shows the minimum read count required in both paired estimates, and the y-axis shows Pearson r of $S_{t3/t0}$. Points represent individual pairwise comparisons, lines show the mean correlation, and shaded regions indicate the SEM. The dashed vertical line marks the gyōza[1] confidence threshold of 50 reads; the dotted vertical line marks the experimental target coverage of 500 reads per variant per library per barcode. Correlations generally improve after low-count variants are removed and a plateau is reached near the intended coverage range, supporting robust downstream interpretation of replicated high-confidence effects.

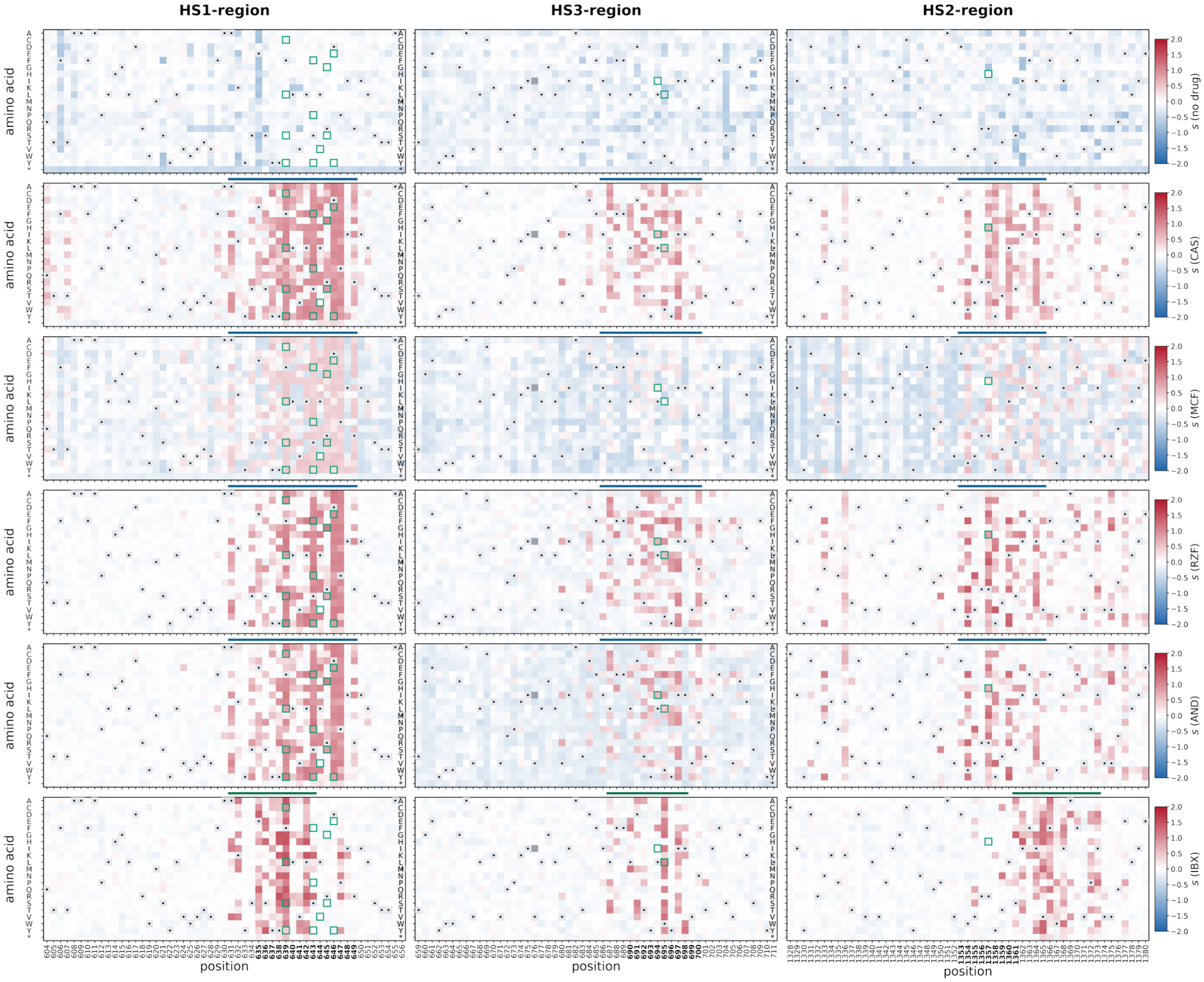


**Figure S6. Full mutation-level fitness landscapes across the complete DMS-targeted *FKS1* regions.** Nested mean selection coefficients (S) are shown for every single amino acid substitution across the full HS1-, HS3-, and HS2-targeted DMS regions. Rows show no drug, CAS, MCF, RZF, AND, and IBX. Red indicates enrichment relative to the matched $t_0$ input population, whereas blue indicates depletion. Wild-type residues are shown as black dots on light grey tiles, and unscored substitutions are shown as dark grey tiles. Green outlines mark priority substitutions identified from recurrent *C. auris* population variation in the Gifford et al.[2] dataset and from the most frequently reported echinocandin-resistance-associated substitutions in FungAMR[3], shown in **Supplementary Table S2**. Historical hotspot positions are indicated by bold black residue numbers. This figure provides the complete regional context for the cropped mutation-level heatmaps shown in **Figure 2A**.

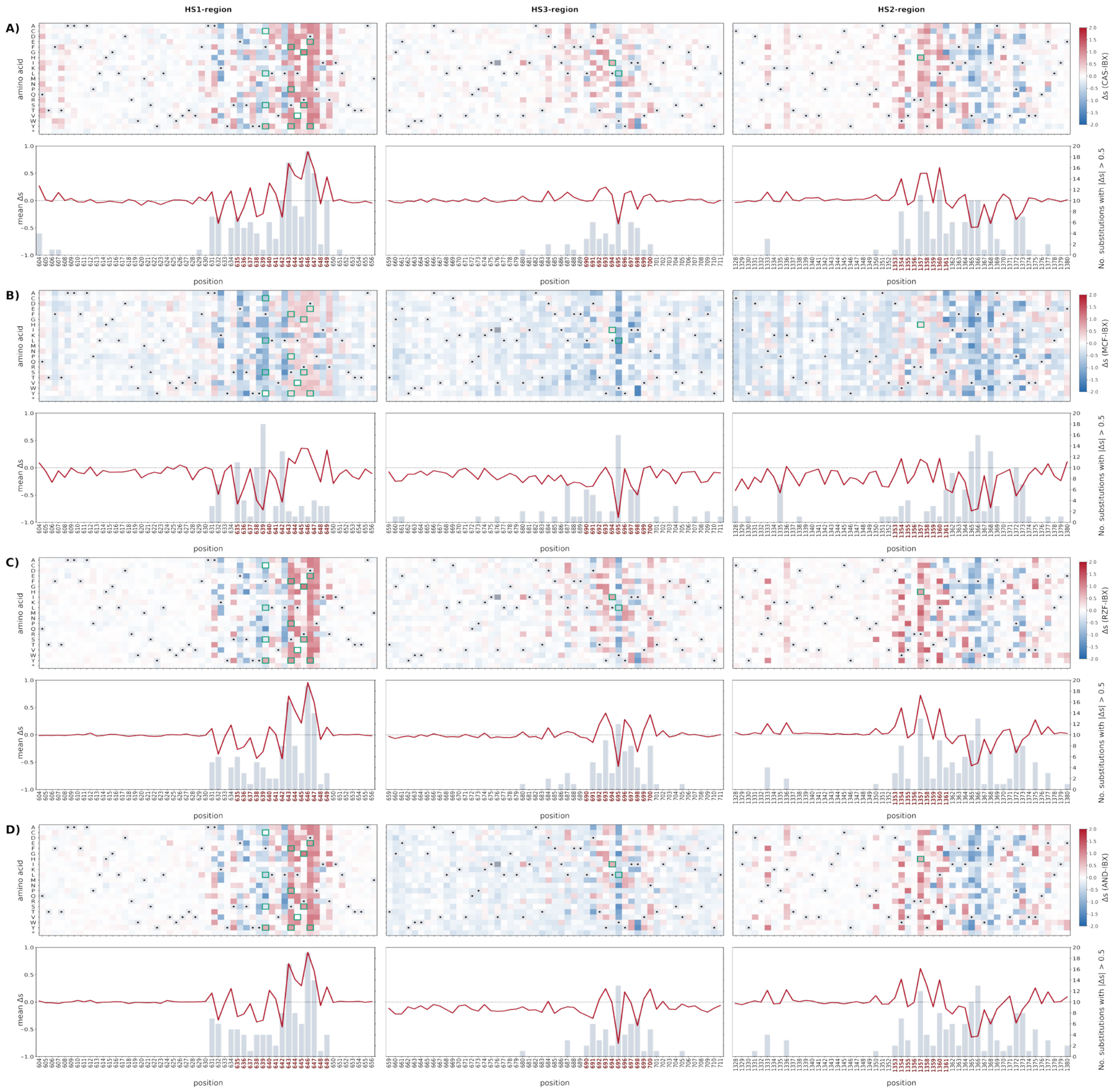


**Figure S7. Differential mutation-level selection between echinocandins and ibrexafungerp.** Pairwise differential selection maps comparing each echinocandin condition with IBX across the full DMS-targeted regions. For each mutation, differential selection was calculated as $\Delta S_{m,A-IBX} = S_{m,A} - S_{m,IBX}$, where $A$ indicates CAS, MCF, RZF, or AND. Red indicates stronger enrichment under the echinocandin condition, whereas blue indicates stronger enrichment under IBX. Green outlines mark priority substitutions identified from Gifford et al.[2] and FungAMR[3], shown in **Supplementary Table S2**. For each comparison, the plots below the heatmaps summarize position-level divergence: the red line shows the mean differential selection coefficient $(\overline{\Delta S})$, and grey bars show the number of substitutions with $|\Delta S| > 0.5$.

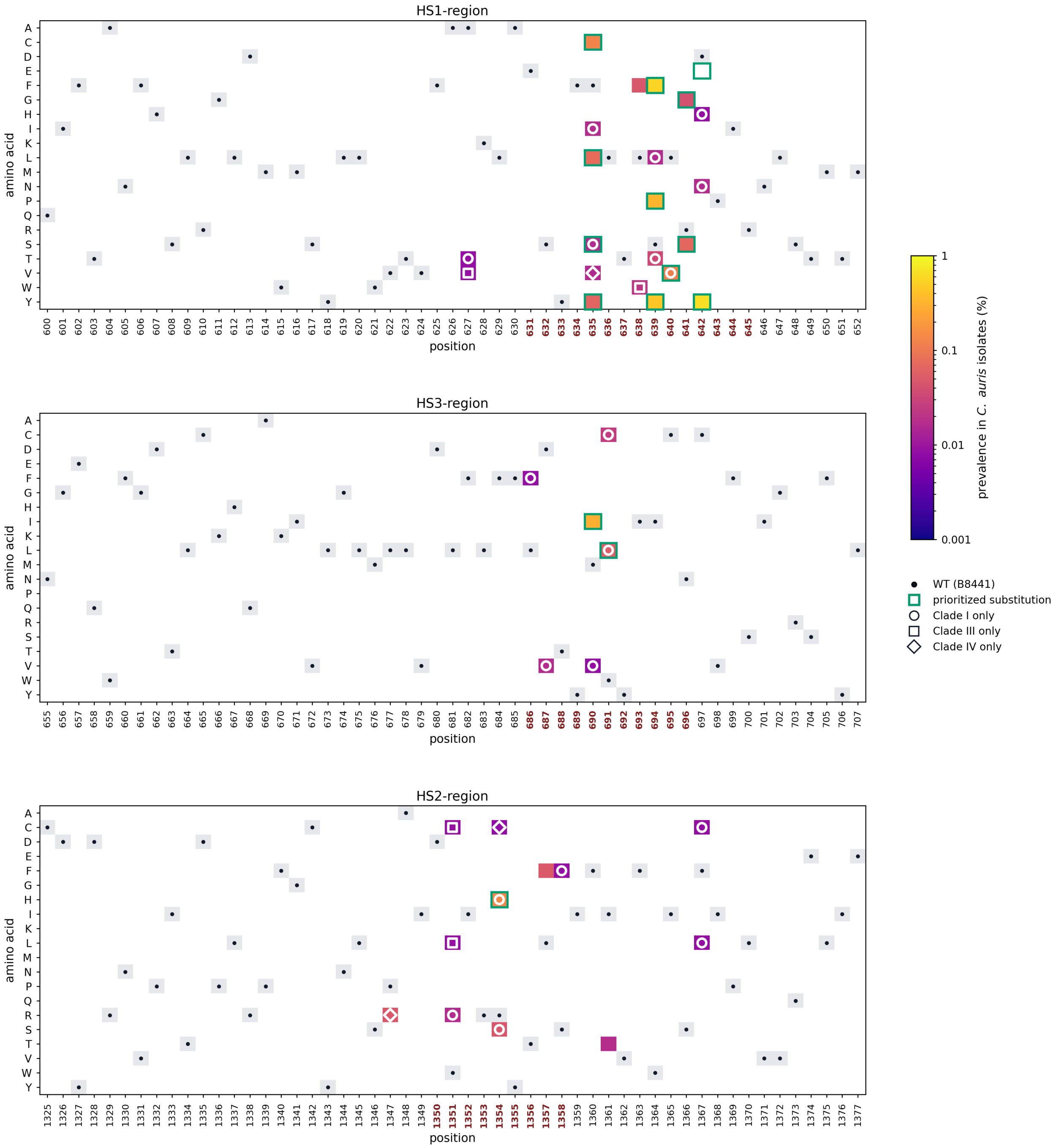


**Figure S8. Prevalence of *FKS1* substitutions in *C. auris* and prioritized resistance-associated substitutions.** Heatmaps show the prevalence of *FKS1* amino acid substitutions in 12,644 *C. auris* isolates from the Gifford et al. dataset[2], restricted to the *C. auris* regions corresponding to the *S. cerevisiae* DMS-targeted HS1-, HS3-, and HS2-regions. Tile colour indicates overall prevalence (%) across the analyzed *C. auris* clades. Grey tiles with black dots indicate the wild-type amino acid in the *C. auris* B8441 reference strain. White symbols indicate substitutions observed in only one clade. Historical hotspot positions, mapped from the corresponding *S. cerevisiae* hotspot intervals, are highlighted in red on the x-axis. Green boxes mark prioritized substitutions, defined as the union of the ten most frequently reported *FKS1* echinocandin-resistance substitutions in FungAMR[3] (see **Supplementary Table S2**) and substitutions present at ≥0.05% prevalence in the Gifford et al. *C. auris* dataset. Deletions, insertions, and stop codons were excluded because the DMS analysis focused on amino acid substitutions.

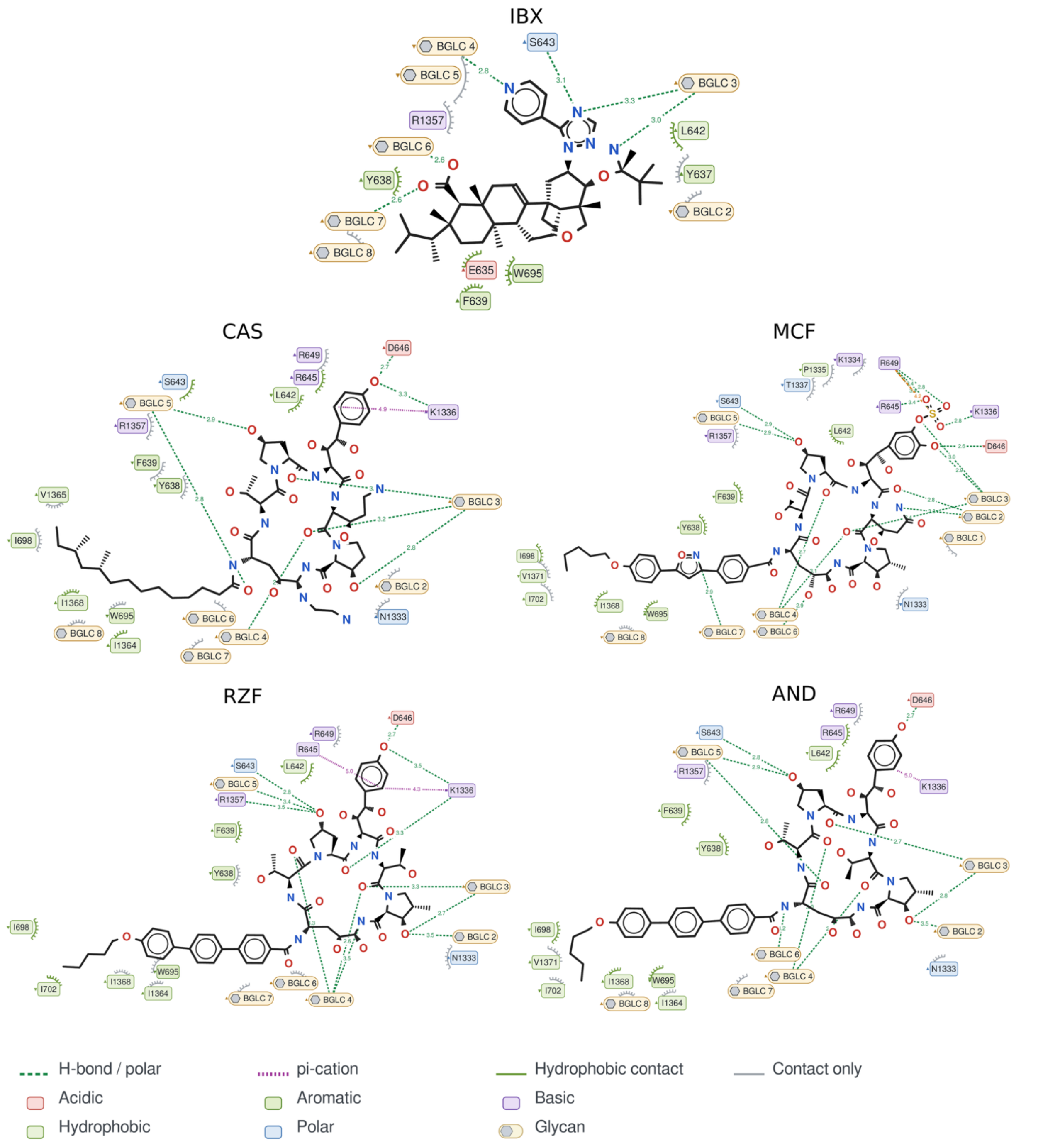


**Figure S9. Protein–ligand interaction maps for IBX, CAS, MCF, RZF, and AND.** Interactions between the protein and each compound were predicted by MD-based simulated annealing. The diagrams show the amino acid residues, glycans (BGLC) and small molecules (ergosterols here, ERG) involved in ligand binding, as well as the types of predicted interactions. Green dashed lines represent hydrogen bonds and polar interactions, magenta dotted lines indicate π-cation interactions, green curves represent hydrophobic contacts, and grey curves indicate contacts without a specific interaction assignment. Contacts are within 5 Å of the protein. Amino acid residues are color-coded according to their chemical properties: acidic (red), hydrophobic (light green), aromatic (green), basic (purple), and polar (blue).

#### Supplementary Tables

**Table S1. Literature-prioritized *FKS1* positions from FungAMR[3].**

The table lists the ten most frequently mutated Fks1 positions associated with echinocandin resistance in FungAMR, mapped to *S. cerevisiae* Fks1 coordinates. Residues are ranked by the number of unique FungAMR source identifiers reporting echinocandin-resistance-associated substitutions mapped to the corresponding *S. cerevisiae FKS1* residue. The table reports the mapped *S. cerevisiae* residue and hotspot region, the number of unique supporting reports, the best and complete set of FungAMR confidence scores (C.S.) reported for substitutions at that residue, the number and names of species in which substitutions were reported, the echinocandin-class drugs represented, and the reported substitutions mapped to the *S. cerevisiae* Fks1 sequence. FungAMR confidence scores grade the type and strength of evidence supporting a mutation-resistance association; lower positive scores indicate stronger evidence. Score 1 denotes experimental introduction of the mutation into the same susceptible species background with phenotypic confirmation; score 2 denotes experimental validation in a non-native species or expression context; score 3 denotes evidence from bulk competition or deep-mutational-scanning-type assays; score 4 denotes recurrent enrichment in experimental evolution; scores 5 and 6 denote resistance linked to gene deletion or gene overexpression/duplication, respectively; score 7 denotes population-level association such as GWAS, QTL, or classical genetics; and score 8 denotes observation in a resistant natural or clinical isolate without further validation. Abbreviations: ECN, echinocandin-class entry; AND, anidulafungin; CAS, caspofungin; MCF, micafungin; RZF, rezafungin/CD101.

| Rank | ScFks1 residue | Region | Unique reports | Best C.S. | C.S. reported | Species (n) | Species | ECN drugs reported | Mapped *S. cerevisiae* substitutions |
|---|---|---|---|---|---|---|---|---|---|
| **1** | S643 | HS1 | 38 | 1 | 1; 2; 3; 4; 7; 8 | 11 | *C. albicans; C. dubliniensis; C. metapsilosis; C. parapsilosis; C. tropicalis; C. auris; C. lusitaniae; K. marxianus; N. glabratus; P. kudriavzevii; S. cerevisiae* | AND; CD101/RZF; CAS; ECN; MCF | S643C; S643D; S643E; S643F; S643G; S643H; S643I; S643K; S643L; S643M; S643N; S643P; S643Q; S643R; S643S; S643T; S643V; S643W; S643X; S643Y |
| **2** | F639 | HS1 | 23 | 1 | 1; 2; 3; 4; 7; 8 | 8 | *C. albicans; C. tropicalis; C. auris; C. lusitaniae; K. marxianus; N. glabratus; P. kudriavzevii; S. cerevisiae* | AND; CAS; ECN; MCF | F639A; F639C; F639D; F639DEL; F639E; F639G; F639H; F639I; F639K; F639L; F639N; F639P; F639Q; F639R; F639S; F639T; F639V; F639W; F639Y |
| **3** | D646 | HS1 | 15 | 1 | 1; 3; 7; 8 | 4 | *C. albicans; C. auris; N. glabratus; S. cerevisiae* | AND; CAS; ECN; MCF | D646A; D646C; D646E; D646F; D646G; D646H; D646I; D646K; D646L; D646M; D646N; D646P; D646Q; D646R; D646S; D646T; D646V; D646W; D646X; D646Y |
| **4** | R645 | HS1 | 10 | 1 | 1; 2; 3; 8 | 4 | *C. albicans; K. marxianus; N. glabratus; S. cerevisiae* | AND; CAS; ECN; MCF | R645A; R645C; R645D; R645E; R645F; R645G; R645H; R645M; R645N; R645P; R645Q; R645S; R645T; R645W; R645Y |
| **5** | R1357 | HS2 | 7 | 1 | 1; 3; 8 | 5 | *C. albicans; C. auris; C. lusitaniae; P. kudriavzevii; S. cerevisiae* | AND; CAS; MCF | R1357C; R1357F; R1357G; R1357H; R1357I; R1357L; R1357M; R1357N; R1357P; R1357Q; R1357S; R1357T; R1357V; R1357W; R1357Y |
| **6** | P647 | HS1 | 6 | 1 | 1; 2; 3; 7; 8 | 3 | *C. albicans; C. lusitaniae; S. cerevisiae* | AND; CAS; MCF | P647A; P647D; P647E; P647F; P647G; P647H; P647I; P647K; P647L; P647M; P647N; P647Q; P647R; P647S; P647T; P647V; P647Y |
| **7** | L644 | HS1 | 4 | 3 | 3; 7; 8 | 3 | *C. tropicalis; N. glabratus; S. cerevisiae* | AND; CAS; MCF | L644E; L644H; L644I; L644K; L644Q; L644R; L644W |
| **8** | W1354 | HS2 | 3 | 3 | 3; 7; 8 | 3 | *C. albicans; C. parapsilosis; S. cerevisiae* | AND; CAS; MCF | W1354F; W1354G; W1354H; W1354L; W1354M; W1354P; W1354R; W1354T; W1354Y |
| **9** | W695 | HS3 | 2 | 1 | 1; 8 | 2 | *C. auris; S. cerevisiae* | AND; CAS; MCF | W695C; W695F; W695L |
| **10** | L690 | HS3 | 2 | 1 | 1; 8 | 2 | *C. parapsilosis; S. cerevisiae* | AND; CAS; MCF | L690A; L690E; L690F |

**Table S2. Literature-prioritized *FKS1* substitutions from FungAMR**[3]. The table lists the ten most frequently reported Fks1 substitutions associated with echinocandin resistance in FungAMR mapped to *S. cerevisiae* Fks1 coordinates. Ranking was based on the number of independent FungAMR source identifiers supporting each exact substitution; ties were resolved by confidence score (C.S.), number of species, and total number of supporting entries. The FungAMR confidence score is shown as reported in the database, with lower scores indicating stronger evidence. Confidence score definitions and abbreviations are given in the caption of **Table S1**.

| Rank | ScFks1 substitution | Region | Unique reports | Best C.S. | C.S. reported | Species (n) | Species reporting substitution | ECN drugs reported |
|---|---|---|---|---|---|---|---|---|
| **1** | S643P | HS1 | 33 | 1 | 1, 2, 4, 7, 8 | 11 | *C. albicans; C. dubliniensis; C. metapsilosis; C. parapsilosis; C. tropicalis; C. auris; C. lusitaniae; K. marxianus; N. glabratus; P. kudriavzevii; S. cerevisiae* | AND, CD101, CAS, ECN, MCF |
| **2** | F639S | HS1 | 16 | 1 | 1, 4, 7, 8 | 5 | *C. albicans; C. tropicalis; K. marxianus; N. glabratus; S. cerevisiae* | AND, CAS, ECN, MCF |
| **3** | S643F | HS1 | 12 | 1 | 1, 4, 7, 8 | 5 | *C. albicans; C. auris; N. glabratus; P. kudriavzevii; S. cerevisiae* | AND, CAS, MCF |
| **4** | S643Y | HS1 | 9 | 1 | 1, 4, 7, 8 | 4 | *C. albicans; C. auris; N. glabratus; S. cerevisiae* | AND, CAS, MCF |
| **5** | D646Y | HS1 | 8 | 3 | 3, 7, 8 | 4 | *C. albicans; C. auris; N. glabratus; S. cerevisiae* | AND, CAS, MCF |
| **6** | D646E | HS1 | 8 | 3 | 3, 7, 8 | 2 | *N. glabratus; S. cerevisiae* | AND, CAS, ECN, MCF |
| **7** | R645G | HS1 | 7 | 1 | 1, 2, 3, 8 | 4 | *C. albicans; K. marxianus; N. glabratus; S. cerevisiae* | AND, CAS, ECN, MCF |
| **8** | F639Y | HS1 | 5 | 2 | 2, 3, 8 | 3 | *C. auris; N. glabratus; S. cerevisiae* | AND, CAS, MCF |
| **9** | F639C | HS1 | 4 | 1 | 1, 4, 8 | 4 | *C. albicans; C. auris; P. kudriavzevii; S. cerevisiae* | AND, CAS, MCF |
| **10** | R1357H | HS2 | 4 | 1 | 1, 3, 8 | 3 | *C. albicans; C. auris; S. cerevisiae* | AND, CAS, MCF |

**Table S3. Information on all strains used in this study.**

| name | species | Fks1 | information | origin |
|---|---|---|---|---|
| R1158 | *Saccharomyces cerevisiae* | WT | wild-type reference | 4 |
| HC001 | *Saccharomyces cerevisiae* | HS1::HPH | constructed mutant from R1158 | this study |
| HC002 | *Saccharomyces cerevisiae* | HS2::HPH | constructed mutant from R1158 | this study |
| HC003 | *Saccharomyces cerevisiae* | HS3::HPH | constructed mutant from R1158 | this study |
| BY4741 | *Saccharomyces cerevisiae* | WT | wild-type reference | 5 |
| s015_1 | *Saccharomyces cerevisiae* | F639C | constructed mutant from BY4741 | 6 |
| s042_24 | *Saccharomyces cerevisiae* | F639Q | constructed mutant from BY4741 | 6 |
| s028_34 | *Saccharomyces cerevisiae* | L640K | constructed mutant from BY4741 | 6 |
| s029_36 | *Saccharomyces cerevisiae* | V641S | constructed mutant from BY4741 | 6 |
| s17_30 | *Saccharomyces cerevisiae* | L642G | constructed mutant from BY4741 | 6 |
| s036_44 | *Saccharomyces cerevisiae* | L642K | constructed mutant from BY4741 | 6 |
| s025_10 | *Saccharomyces cerevisiae* | S643F | constructed mutant from BY4741 | 6 |
| s039_21 | *Saccharomyces cerevisiae* | R645S | constructed mutant from BY4741 | 6 |
| s031_15 | *Saccharomyces cerevisiae* | D646W | constructed mutant from BY4741 | 6 |
| s019_3 | *Saccharomyces cerevisiae* | P647N | constructed mutant from BY4741 | 6 |
| B8441 | *Candidozyma auris* | WT | wild-type reference Clade I (Pakistan) | 7 |
| B11220 | *Candidozyma auris* | WT | wild-type reference Clade II (Japan) | 8 |
| NCPF8971 | *Candidozyma auris* | WT | wild-type reference Clade I (UK) | 9 |
| C3 | *Candidozyma auris* | F635Δ | experimentally evolved from B11220 | 10 |
| C15 | *Candidozyma auris* | F635Δ+M690I | experimentally evolved from B11220 | 10 |
| C2_2 | *Candidozyma auris* | F635Y+S639Y | experimentally evolved from B8441 | 11 |
| UACa141 | *Candidozyma auris* | S639P | experimentally evolved from B11220 | 12 |
| UACa142 | *Candidozyma auris* | S639P | experimentally evolved from B11220 | 12 |

| | | | | |
|---|---|---|---|---|
| UACa143 | *Candidozyma auris* | S639Y | experimentally evolved from B11220 | 12 |
| LC736030 | *Candidozyma auris* | S639Y | experimentally evolved from NCPF8971 | 9 |
| LC736029 | *Candidozyma auris* | R641G | experimentally evolved from NCPF8971 | 9 |
| LC736033 | *Candidozyma auris* | R1354H | experimentally evolved from NCPF8971 | 9 |
| LC736032 | *Candidozyma auris* | R1354Y | experimentally evolved from NCPF8971 | 9 |
| 152 | *Candidozyma auris* | S639Y | experimentally evolved from B8441 | 11,13 |
| C10_1 | *Candidozyma auris* | S639P | experimentally evolved from B8441 | 11 |
| F11_1 | *Candidozyma auris* | W691L | experimentally evolved from B8441 | 11 |
| F883 | *Candidozyma auris* | S639P | clinical isolate (Belgium) | 14 |
| NCCPF470238 | *Candidozyma auris* | F635L | clinical isolate (India) | 15 |
| NCCPF470237 | *Candidozyma auris* | F635L | clinical isolate (India) | 15 |
| NCCPF470240 | *Candidozyma auris* | F635L | clinical isolate (India) | 15 |
| NCCPF470241 | *Candidozyma auris* | F635L | clinical isolate (India) | 15 |
| NCCPF470197 | *Candidozyma auris* | F635Y | clinical isolate (India) | 15 |
| NCCPF470199 | *Candidozyma auris* | S639F | clinical isolate (India) | 15 |
| NCCPF470198 | *Candidozyma auris* | S639F | clinical isolate (India) | 15 |
| NCCPF470200 | *Candidozyma auris* | R1354S | clinical isolate (India) | 15 |
| Kw1893/17 | *Candidozyma auris* | R1354H | clinical isolate (Kuwait) | 16 |
| Kw2074/17 | *Candidozyma auris* | R1354H | clinical isolate (Kuwait) | 16 |
| Kw281/18 | *Candidozyma auris* | S639F | clinical isolate (Kuwait) | 16 |
| Kw2742/17 | *Candidozyma auris* | S639F | clinical isolate (Kuwait) | 16 |
| Kw1994/17 | *Candidozyma auris* | S639F | clinical isolate (Kuwait) | 16 |
| Kw108/18 | *Candidozyma auris* | S639T | clinical isolate (Kuwait) | 16 |
| Kw87/18 | *Candidozyma auris* | S639T | clinical isolate (Kuwait) | 16 |
| Kw3134/17 | *Candidozyma auris* | S639P | clinical isolate (Kuwait) | 17 |
| Kw2713/17 | *Candidozyma auris* | S639F | clinical isolate (Kuwait) | 17 |
| Kw1308/17 | *Candidozyma auris* | F635Δ | clinical isolate (Kuwait) | 18 |
| HC004-6 | *Candidozyma auris* | F635Δ | constructed mutant from B8441 | this study |
| HC007-9 | *Candidozyma auris* | F635C | constructed mutant from B8441 | this study |
| HC010-12 | *Candidozyma auris* | F635Y | constructed mutant from B8441 | this study |
| HC013-15 | *Candidozyma auris* | S639F | constructed mutant from B8441 | this study |
| HC016-18 | *Candidozyma auris* | S639P | constructed mutant from B8441 | this study |
| HC019-21 | *Candidozyma auris* | S639Y | constructed mutant from B8441 | this study |
| HC022-24 | *Candidozyma auris* | D642H | constructed mutant from B8441 | this study |
| HC025-27 | *Candidozyma auris* | D642H+R645T | constructed mutant from B8441 | this study |
| HC028-30 | *Candidozyma auris* | M690V | constructed mutant from B8441 | this study |
| HC031-33 | *Candidozyma auris* | F635Δ+M690I | constructed mutant from B8441 | this study |
| HC034-36 | *Candidozyma auris* | W691L | constructed mutant from B8441 | this study |

**Table S4. Plasmids and oligonucleotides used in this study.**

Materials used for strain construction, DMS library preparation and validation, and genotyping are listed with their name, type, use, and sequence or origin. Oligonucleotide sequences are presented in the 5′–3′ direction.

| name | type | use | sequence/origin |
|---|---|---|---|
| pCAS-HPH | plasmid | CRISPR–Cas9 plasmid for *S. cerevisiae FKS1* DMS library integration. | 19 |
| pFA-hphNT1 | plasmid | hphNT1 cassette for *S. cerevisiae* recipient construction. | 20 |
| pJMR19 | plasmid | Backbone for the *C. auris FKS1* editing plasmids pHC26_1–3. | 21 |
| pHC26_1 | plasmid | pJMR19 derivative for construction of *C. auris FKS1* F635 variants. | repository link available upon formal publication |
| pHC26_2 | plasmid | pJMR19 derivative for construction of *C. auris FKS1* S639/D642 variants. | repository link available upon formal publication |
| pHC26_3 | plasmid | pJMR19 derivative for construction of *C. auris FKS1* M690/W691 variants. | repository link available upon formal publication |
| CLOP353-G1 | primer | hphNT1 cassette PCR for *S. cerevisiae FKS1* HS1 recipient construction (forward). | TACGTCATATATGAAAAATCTACAAGGCGTTATGTTGCAtcgtacgctgcaggtcgac |

| | | | |
|---|---|---|---|
| CLOP353-H1 | primer | hphNT1 cassette PCR for *S. cerevisiae FKS1* HS1 recipient construction (reverse). | GCACTTTACAAAGTACCGCACCCCACCAGTATTCACCTGTatcgatgaattcgagctcg |
| CLOP353-C2 | primer | hphNT1 cassette PCR for *S. cerevisiae FKS1* HS2 recipient construction (forward). | GACTTTGGTGAATTTATCTTCCTTGGCCCATGAATCTATTcgtacgctgcaggtcgac |
| CLOP353-D2 | primer | hphNT1 cassette PCR for *S. cerevisiae FKS1* HS2 recipient construction (reverse). | ATAGGTGGCAGAAAAATCTTTGGGTGGCTTTCCATAGACCatcgatgaattcgagctcg |
| CLOP353-A2 | primer | hphNT1 cassette PCR for *S. cerevisiae FKS1* HS3 recipient construction (forward). | TTTGAGAGATCCAATTAGAATTTTGTCCACCACTGCAATGtcgtacgctgcaggtcgac |
| CLOP353-B2 | primer | hphNT1 cassette PCR for *S. cerevisiae FKS1* HS3 recipient construction (reverse). | ATCTTGTGAAGATATTTCTCCATGGTGTTAAGATAGAAATatcgatgaattcgagctcg |
| CLOP353-E2 | primer | *S. cerevisiae FKS1* HS1 recipient verification by colony PCR (upstream flanking primer). | CGTAGATTCTGGTTTTTATGCATCATCTTTGG |
| CLOP353-F2 | primer | *S. cerevisiae FKS1* HS1 recipient verification by colony PCR (downstream flanking primer). | GCCAAAATCTTGGAGTATATTCTTTTTGGC |
| CLOP353-A3 | primer | *S. cerevisiae FKS1* HS2 recipient verification by colony PCR (upstream flanking primer). | CAATTCTAAATTTCACTACTAAGATTGGTGCTGG |
| CLOP353-B3 | primer | *S. cerevisiae FKS1* HS2 recipient verification by colony PCR (downstream flanking primer). | ACCCATGTAGATAGCAGATCCTGC |
| CLOP353-G2 | primer | *S. cerevisiae FKS1* HS3 recipient verification by colony PCR (upstream flanking primer). | TTGCTACCATCATATTCTTCTCCATTATGC |
| CLOP353-H2 | primer | *S. cerevisiae FKS1* HS3 recipient verification by colony PCR (downstream flanking primer). | GGAGCTCTCAAAGTTCTTTTACCTTCG |
| CLO1-38 | primer | *S. cerevisiae* recipient colony PCR: hphNT1-internal primer, 5′ junction. | GTCGCGGTGAGTTCAGGCTT |
| CLOP81-H1 | primer | *S. cerevisiae* recipient colony PCR: hphNT1-internal primer, 3′ junction. | CCGATAGTGGAAACCGACGC |
| CLOP102-G2 | gRNA oligo | Insertion of the hphNT1-targeting gRNA into pCAS (forward). | cgggtggcgaatgggactttgaatcggtcaatacactacagttttagagctagaaatagc |
| CLOP102-H2 | gRNA oligo | Insertion of the hphNT1-targeting gRNA into pCAS (reverse). | gctatttctagctctaaaactgtagtgtattgaccgattcaaagtcccattcgccacccg |
| CLOP366-A12 | primer | HS1 donor-oPool PCR for *S. cerevisiae FKS1* library construction (forward). | TACGTCATATATGAAAAAATCTACAAGGCGTTATGTTGCA |
| CLOP366-B12 | primer | HS1 donor-oPool PCR for *S. cerevisiae FKS1* library construction (reverse). | GCACTTTACAAAGTACCGCACCCCACCAGTATTCACCTGTACA |
| CLOP366-E12 | primer | HS2 donor-oPool PCR for *S. cerevisiae FKS1* library construction (forward). | GACTTTGGTGAATTTATCTTCCTTGGCCCATGAATCTATT |
| CLOP366-F12 | primer | HS2 donor-oPool PCR for *S. cerevisiae FKS1* library construction (reverse). | ATAGGTGGCAGAAAAATCTTTGGGTGGCTTTCCATAGACC |
| CLOP366-C12 | primer | HS3 donor-oPool PCR for *S. cerevisiae FKS1* library construction (forward). | TTTGAGAGATCCAATTAGAATTTTGTCCACCACTGCAATGAGG |
| CLOP366-D12 | primer | HS3 donor-oPool PCR for *S. cerevisiae FKS1* library construction (reverse). | ATCTTGTGAAGATATTTCTCCATGGTGTTAAGATAGAAAT |
| CLOP372-A8 | primer | *S. cerevisiae FKS1* HS1 $t_0/t_3$ amplicon-sequencing PCR (LIB1, forward). | TCGTCGGCAGCGTCAGATGTGTATAAGAGACAGNAATCTACAAGGCGTTATGTTGC |
| CLOP372-H8 | primer | *S. cerevisiae FKS1* HS1 $t_0/t_3$ amplicon-sequencing PCR (LIB1, reverse). | GTCTCGTGGGCTCGGAGATGTGTATAAGAGACAGNNCACCCCACCAGTATTCACC |
| CLOP372-B8 | primer | *S. cerevisiae FKS1* HS1 $t_0/t_3$ amplicon-sequencing PCR (LIB2, forward). | TCGTCGGCAGCGTCAGATGTGTATAAGAGACAGNNAATCTACAAGGCGTTATGTTGC |
| CLOP372-G8 | primer | *S. cerevisiae FKS1* HS1 $t_0/t_3$ amplicon-sequencing PCR (LIB2, reverse). | GTCTCGTGGGCTCGGAGATGTGTATAAGAGACAGNCACCCCACCAGTATTCACC |
| CLOP372-C8 | primer | *S. cerevisiae FKS1* HS2 $t_0/t_3$ amplicon-sequencing PCR (LIB1, forward). | TCGTCGGCAGCGTCAGATGTGTATAAGAGACAGNGGTGAATTTATCTTCCTTGGC |
| CLOP372-B9 | primer | *S. cerevisiae FKS1* HS2 $t_0/t_3$ amplicon-sequencing PCR (LIB1, reverse). | GTCTCGTGGGCTCGGAGATGTGTATAAGAGACAGNNGGGTGGCTTTCCATAGACC |
| CLOP372-D8 | primer | *S. cerevisiae FKS1* HS2 $t_0/t_3$ amplicon-sequencing PCR (LIB2, forward). | TCGTCGGCAGCGTCAGATGTGTATAAGAGACAGNNGGTGAATTTATCTTCCTTGGC |
| CLOP372-A9 | primer | *S. cerevisiae FKS1* HS2 $t_0/t_3$ amplicon-sequencing PCR (LIB2, reverse). | GTCTCGTGGGCTCGGAGATGTGTATAAGAGACAGNGGGTGGCTTTCCATAGACC |
| CLOP372-E8 | primer | *S. cerevisiae FKS1* HS3 $t_0/t_3$ amplicon-sequencing PCR (LIB1, forward). | TCGTCGGCAGCGTCAGATGTGTATAAGAGACAGNGAATTTTGTCCACCACTGC |
| CLOP372-D9 | primer | *S. cerevisiae FKS1* HS3 $t_0/t_3$ amplicon-sequencing PCR (LIB1, reverse). | GTCTCGTGGGCTCGGAGATGTGTATAAGAGACAGNNGTGAAGATATTTCTCCATGGTG |
| CLOP372-F8 | primer | *S. cerevisiae FKS1* HS3 $t_0/t_3$ amplicon-sequencing PCR (LIB2, forward). | TCGTCGGCAGCGTCAGATGTGTATAAGAGACAGNNGAATTTTGTCCACCACTGC |

| | | | |
|---|---|---|---|
| CLOP372-C9 | primer | *S. cerevisiae FKS1* HS3 $t_0$/$t_3$ amplicon-sequencing PCR (LIB2, reverse). | GTCTCGTGGGCTCGGAGATGTGTATAAGAGACAGNGTGAAGATATTTCTCCATGGTG |
| CLOP366-E10 | primer | *S. cerevisiae* HS1 library-validation sequencing: coverage and variant representation (forward 1). | TCGTCGGCAGCGTCAGATGTGTATAAGAGACAGNCTTCTCCATTATGCCATTGG |
| CLOP366-F10 | primer | *S. cerevisiae* HS1 library-validation sequencing: coverage and variant representation (forward 2). | TCGTCGGCAGCGTCAGATGTGTATAAGAGACAGNNCTTCTCCATTATGCCATTGG |
| CLOP366-G10 | primer | *S. cerevisiae* HS1 library-validation sequencing: coverage and variant representation (reverse 1). | GTCTCGTGGGCTCGGAGATGTGTATAAGAGACAGNCAAACCTAAGACAATCTTGG |
| CLOP366-H10 | primer | *S. cerevisiae* HS1 library-validation sequencing: coverage and variant representation (reverse 2). | GTCTCGTGGGCTCGGAGATGTGTATAAGAGACAGNNCAAACCTAAGACAATCTTGG |
| CLOP372-C7 | primer | *S. cerevisiae* HS2 library-validation sequencing: coverage and variant representation (forward 1). | TCGTCGGCAGCGTCAGATGTGTATAAGAGACAGNAATTATCTTTGCAAATGTTTATG |
| CLOP372-D7 | primer | *S. cerevisiae* HS2 library-validation sequencing: coverage and variant representation (forward 2). | TCGTCGGCAGCGTCAGATGTGTATAAGAGACAGNNAATTATCTTTGCAAATGTTTATG |
| CLOP372-G7 | primer | *S. cerevisiae* HS2 library-validation sequencing: coverage and variant representation (reverse 1). | GTCTCGTGGGCTCGGAGATGTGTATAAGAGACAGNGCAAACACTTCGAACATAG |
| CLOP372-H7 | primer | *S. cerevisiae* HS2 library-validation sequencing: coverage and variant representation (reverse 2). | GTCTCGTGGGCTCGGAGATGTGTATAAGAGACAGNNGCAAACACTTCGAACATAG |
| CLOP366-A11 | primer | *S. cerevisiae* HS3 library-validation sequencing: coverage and variant representation (forward 1). | TCGTCGGCAGCGTCAGATGTGTATAAGAGACAGNCTGCCAAATATTCAGAATCG |
| CLOP366-B11 | primer | *S. cerevisiae* HS3 library-validation sequencing: coverage and variant representation (forward 2). | TCGTCGGCAGCGTCAGATGTGTATAAGAGACAGNNCTGCCAAATATTCAGAATCG |
| CLOP366-C11 | primer | *S. cerevisiae* HS3 library-validation sequencing: coverage and variant representation (reverse 1). | GTCTCGTGGGCTCGGAGATGTGTATAAGAGACAGNTGGAGTATATTCTTTTTGGC |
| CLOP366-D11 | primer | *S. cerevisiae* HS3 library-validation sequencing: coverage and variant representation (reverse 2). | GTCTCGTGGGCTCGGAGATGTGTATAAGAGACAGNNTGGAGTATATTCTTTTTGGC |
| D-8127 | gRNA oligo | gRNA oligo for construction of *C. auris FKS1* F635 variants. | CCAAGTCAAGAAGAAGTAAGATT |
| D-8128 | gRNA oligo | gRNA oligo for construction of *C. auris FKS1* F635 variants. | AACAATCTTACTTCTTCTTGACT |
| D-8121 | gRNA oligo | gRNA oligo for construction of *C. auris FKS1* S639 and D642 variants. | CCAGTTTCTAATAGGATCTCTCA |
| D-8122 | gRNA oligo | gRNA oligo for construction of *C. auris FKS1* S639 and D642 variants. | AACTGAGAGATCCTATTAGAAAC |
| D-8123 | gRNA oligo | gRNA oligo for construction of *C. auris FKS1* M690 and W691 variants. | CCATCTTCTTGGATACTTACATG |
| D-8124 | gRNA oligo | gRNA oligo for construction of *C. auris FKS1* M690 and W691 variants. | AACCATGTAAGTATCCAAGAAGA |
| D-8571 | primer | *C. auris FKS1* HS1/HS3 donor-DNA PCR from a gBlock (forward). | TTACATGAACAGAAGATCAAG |
| D-8570 | primer | *C. auris FKS1* HS1/HS3 donor-DNA PCR from a gBlock (reverse). | GCCAAGATCTTGGAGTAG |
| D-2964 | primer | *C. auris FKS1* HS1/HS3 allele verification by PCR and Sanger sequencing (forward). | CTGCGAAATCAACACCTTTG |
| D-2965 | primer | *C. auris FKS1* HS1/HS3 allele verification by PCR and Sanger sequencing (reverse). | TTCTTCGCTATCATGCCTTT |
| D-5412 | primer | *C. auris FKS1* HS2 genotyping of evolved/clinical strains by PCR and Sanger sequencing (forward). | AGCAGATGTTGTCCAGAGAA |
| D-5413 | primer | *C. auris FKS1* HS2 genotyping of evolved/clinical strains by PCR and Sanger sequencing (reverse). | AGATTGAGAACGGAATACGAGAA |
